# Insulin Receptor Loss Impairs Mammary Tumorigenesis in Mice

**DOI:** 10.1101/2022.05.06.490906

**Authors:** Lauren Podmore, Yekaterina Poloz, Catherine Iorio, Samar Mouaaz, Kevin Nixon, Petr Smirnov, Brianna McDonnell, Sonya Lam, Bowen Zhang, Pirashaanthy Tharmapalan, Soumili Sarkar, Foram Vyas, Marguerite Ennis, Ryan Dowling, Vuk Stambolic

**Affiliations:** Department of Medical Biophysics, University of Toronto, Princess Margaret Cancer Research Tower, Toronto, Ontario, Canada M5G 1L7; Princess Margaret Cancer Centre, University Health Network, Princess Margaret Cancer Research Tower, Toronto, Ontario, Canada M5G 1L7; F. Hoffmann-La Roche Ltd, Mississauga, Ontario, Canada L5N 5M8.; Applied Statistician, Markham, Ontario, Canada, L3R 6H9

**Keywords:** insulin receptor, BC, mammary tumourigenesis, genetically engineered mouse model, bioenergetic fitness

## Abstract

Breast cancer (BC) prognosis and outcome are adversely affected by obesity. Hyperinsulinemia, common in obese state, is associated with higher risk of death and recurrence in BC. Up to 80% of breast cancers overexpress the insulin receptor (INSR), which correlates with worse prognosis. INSR role in mammary tumorigenesis was tested by generating MMTV-driven polyoma middle T (PyMT) and ErbB2/Her2 BC mouse models, respectively, with coordinate mammary epithelium-restricted deletion of *INSR*. In both models, deletion of either one or both copies of *INSR* led to a marked delay in tumor onset and burden. Longitudinal phenotypic characterization of mouse tumours and cells revealed that INSR deletion impacted tumour initiation, not progression and metastasis. INSR upheld a bioenergetic phenotype in non-transformed mammary epithelial cells, independent of its kinase activity. Similarity of phenotypes elicited by deletion of one or both copies of *INSR* suggest a dose-dependent threshold for INSR impact on mammary tumorigenesis.

## Introduction

Obesity is a global epidemic, with over 650 million obese and further 1.3 billion overweight individuals worldwide ^1^. 5% and 10% of all US cancer cases in men and women, respectively, can be attributed to excess body weight ^2^. Estimates from two decades ago attribute obesity to approximately 14% of the deaths from cancer in men and 20% of the deaths from cancer in women ^3^. In breast cancer (BC), for each 5 kg/m^2^ increase in body mass index (BMI), risk of BC–specific mortality increased by 17-29% ^4^. Importantly, sustained weight loss is associated with lower BC risk in women 50 or older, indicating that the effects of excess body weight on BC are reversible ^5^.

Elevated levels of circulating insulin, or hyperinsulinemia, a condition that parallels obesity ^6^, has an adverse effect on a patient’s prognosis and risk in operable BC ^7–10^. Further, there is a positive association between cancer incidence and insulin dose in persons with type 2 diabetes treated by either human insulin or synthetic insulin analogues ^11^. Adverse effects of insulin and benefits of insulin deprivation in a rat model of mammary tumorigenesis were observed half a century ago ^12,13^. More recently, excess insulin was found to accelerate mammary tumorigenesis in mouse models, whereas insulin reducing agents counter it ^14,15^. Importantly, in a pancreatic cancer mouse model, hyperinsulinemia accelerated tumorigenesis, whereas genetically-mediated insulin reduction decelerated tumorigenesis, separating the impact of elevated insulin on tumor growth from the accompanying hyperglycemia seen in metabolic disease ^16^.

Clinical studies have revealed that 48-92% of breast tumours express insulin’s preferred cell surface receptor, the insulin receptor (INSR) ^17–19^, while others have shown that breast cancers overexpress INSR protein relative to normal breast tissue ^20,21^. INSR is a heterotetrameric receptor tyrosine kinase typically found in muscle, fat, and liver cells, where it propagates metabolic signalling ^22^. There is evidence from a number of mammalian cell lines on the regulation of INSR transcription, including involvement of hormonal and environmental factors (reviewed in ^23^). While metabolic tissues express the INSR-B isoform, cancers predominantly express INSR-A, the foetal INSR isoform capable of mediating mitogenic signaling ^24^. Engagement of insulin with INSR results in activation of the PI3K-Akt signaling pathway, an anabolic cascade central to the control of glucose transport across cell membranes, glycolysis, lipolysis, mRNA translation and cell growth, survival, and migration ^25^. Hyperactivation of PI3K- Akt signaling is found in many human tumours, largely owing to the activating mutations in PIK3CA, the catalytic subunit of PI3K, and loss of function of the tumor suppressor PTEN, which counters PI3K signaling via its lipid phosphatase activity against phosphatidylinositol second messengers produced by PI3K ^26^. Moreover, insulin stimulation of INSR-A also activates the pro-mitogenic Ras-MAPK signaling cascade ^27^.

Preclinical studies in mouse models of type 2 diabetes mellitus (T2DM) have shown that hyperinsulinemia promoted breast cancer growth and metastasis through activation of INSR and its receptor tyrosine kinase family member insulin-like growth factor receptor 1 (IGF1R), specifically through hyperactivation of the PI3K-Akt and Ras-MAPK pathways ^15,28^. IGF1R shares structural homology, ligands, and downstream signaling effectors with INSR, and is similarly overexpressed in human breast cancer ^29^. Moreover, INSR and IGF1R have been found to form hybrid receptors with unique affinities for binding the ligands insulin, IGF1, and IGF2 ^30^, and activation of these hybrid receptors may promote tumourigenesis ^31^. Finally, it is thought that increased insulin levels found in hyperinsulinemic individuals might be capable of overcoming IGF1R’s lower affinity for binding insulin and activating this pro-mitogenic receptor ^15^.

Despite their distinct biological impact in normal tissues, the individual contributions of INSR and IGF1R to insulin-mediated mammary tumourigenesis remain poorly understood ^32^. Owing to the fact that the detection method for the activated state of INSR and IGF1R is based on the recognition of a shared amino acid sequence within their respective tyrosine kinase domains, their activation cannot be uncoupled *in situ* ^33^. Nevertheless, clinical studies have found that increased INSR expression and activation of INSR and/or IGF1R, but not increased IGF1R expression alone, associated with enhanced tumourigenesis and worse survival outcomes in human breast cancer ^18,34^. Given that the mitogenic INSR-A is the predominantly expressed INSR isoform in cancer, these data indicate that INSR alone may be capable of conferring the pro-tumourigenic effects of hyperinsulinemia.

To better understand how INSR individually contributes to the function of insulin signaling in mammary tumorigenesis, we genetically disrupted the *INSR* gene in mammary tissues of two mouse models of BC, driven by the polyoma middle T (*PyMT*) and *HER2/ErbB2/neu* oncogenes, respectively. Our data demonstrate that *INSR* loss impairs mammary tumor initiation, but not progression or metastasis, and suggest the existence of a dose-dependent threshold for the contribution of INSR to mammary tumorigenesis. *INSR* loss in non-transformed murine mammary epithelial cells lowered mRNA expression of many components of the mitochondrial respiratory chain and impairs their energetic capacity, a phenotype rescued by both wt and kinase-dead INSR, implicating bioenergetic fitness as a mechanism by which INSR mediates breast tumour initiation, likely through non-canonical INSR signaling.

## Results

### *INSR* is broadly expressed in human cancer and regulates PI3K-Akt and Ras-MAPK signaling pathways in mammary epithelial cells

Widespread INSR protein presence and preferential mitogenic INSR-A isoform expression in human BC were discovered in the 1990s (**Figure 1A**) ^21^. Subsequent immunohistochemical analyses of human BC revealed that 72-88% of tumour specimens feature high levels of INSR protein (**Figure 1A**) ^17,19–21,35^. More recent proteomic analyses of human BC specimens corroborate prevalent increased INSR levels compared to adjacent normal tissue and lack of association with a particular molecular subtype (**Supplementary Figure 1A-D**)^36,38^. Consistently, broad *INSR* mRNA expression in the Cancer Cell Line Encyclopedia BC dataset ^37^ appeared equal among BC cell lines grouped by molecular subtype (**Supplementry Figure 1E-F**), as classified by the Cancer Dependency Map Project at the Broad Institute (DepMap.org) ^39^. We analyzed *INSR* isoform mRNA expression by RT-qPCR within a select group of cell lines (**Figure 1B**). All cell lines expressed both *INSR* isoforms, but at differing ratios of *INSR-A:INSR-B*, respectively; 40:60 in untransformed mammary epithelial MCF-10A cells, 80:20 in breast cancer cell lines MCF-7 and MDA-MB-231, and 60:40 in kidney cancer-derived HEK-293T cells. On protein level, INSR abundance in MCF-7 and MDA-MB-231 was comparable to that of HEK-293T cells, whereas MCF-10A cells displayed relatively lower INSR expression (**Figure 1C**).

**Figure 1:**
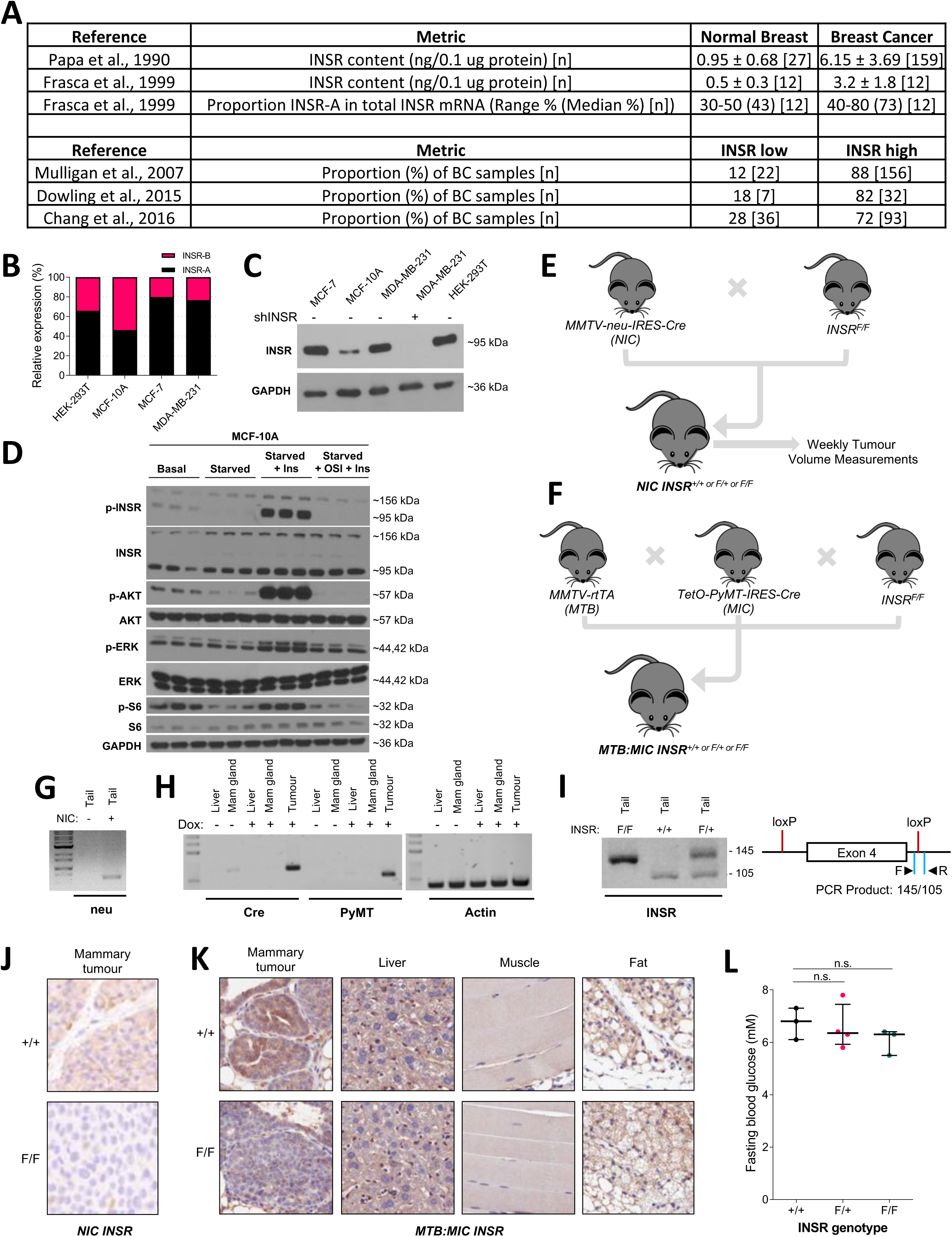
INSR expression in human breast cancer. (**A**)Table summarizing INSR expression data from previous clinical studies. (**B**) Bar graph showing RT-qPCR analysis of relative mRNA expression of *INSR-A* and *INSR-B* isoforms in HEK-293T, MCF-10A, MCF-7 and MDA-MB-231 cell lines. (**C**) Western blot analysis of INSR protein expression in HEK-293T, MCF-7 and MDA-MB-231 and MCF- 10A cell lines. MDA-MB-231 cells were transduced with an shRNA targeting *INSR* to demonstrate specificity. Blots were probed for INSR, with GAPDH probed for as a loading control. (**D**) Western blot analysis of MCF-10A cells grown in supplemented DMEM/HAM (Basal) or starved overnight in the absence of insulin and FBS (Starved), and then either stimulated with 100 nM insulin for 5 minutes (Starved+Ins) or pre-treated with 1 µM OSI-906 for 2 hours before stimulation with 100 nM insulin for 5 minutes (Starved+OSI+Ins). Blots were probed to determine both total and phosphorylated levels of INSR, AKT, ERK and S6. GAPDH was probed for a loading control. Boxplots show the median and interquartile range, with Tukey style whiskers. Scatterplot shows the median and interquartile range. ** p < 0.01, *** p < 0.001, **** p < 0.0001; Student’s t-test. (**E**) Breeding schematic for *NIC* mice that are either wild-type (+/+), heterozygous (F/+) or homozygous (F/F) for the conditional *INSR* allele. Tumour measurements were performed weekly, and mice were sacrificed at humane endpoint. (**F**) Breeding schematic for *MTB:MIC* mice that are either wild-type (+/+), heterozygous (F/+) or homozygous (F/F) for the conditional *INSR* allele. 10-week-old mice were fed either regular fat (RFD) or high fat (HFD) diets for an additional 10 weeks, followed by treatment with 2 mg/ml doxycycline to induce mammary tissue-specific expression of the PyMT oncogene with coordinate deletion of *INSR*. (**G**) Agarose gel image of PCR showing the genotyping of *NIC* positive (+) and negative (-) mice. (**H**) Agarose gel image of PCR showing tissue-specific Cre and PyMT expression upon doxycycline administration in the mammary tumours of *MTB:MIC* mice. (**I**) Agarose gel showing *INSR* exon 4 excision in *INSR^F/+^* (F/+) and *INSR^F/F^* (F/F) mice, followed by the schematic of the PCR strategy. Mice that are wild-type for *INSR* (+/+) are included as a control. (**J**) Immunohistochemical analysis of INSR expression in the mammary gland of *NIC* mice that are either wild-type (+/+) or homozygous (F/F) for the conditional *INSR* allele. (**K**) Immunohistochemical analysis of INSR expression in mammary gland, liver, muscle and fat tissues of *MTB:MIC* mice that are either wild-type (+/+) or homozygous (F/F) for the conditional *INSR* allele. (**L**) Scatterplot of fasting blood glucose levels in *FVB* (+/+, n = 3), *MTB:MIC INSR* (F/+) (n = 4) and *MTB:MIC INSR* (F/F) (n = 3) model mice. Scatterplot shows the median and interquartile range. n.s = non-significant; Student’s t- test.

To probe whether expressed INSR is responsive to ligand stimulation in mammary epithelia, MCF-10A cells were starved of serum and growth factors and short-term stimulated with insulin. Judging by stimulation-dependent increase in phosphorylation of INSR, AKT, S6 and ERK, insulin led to activation of both the metabolic and mitogenic arms of INSR signaling, reflective of their INSR-A expression (**Figure 1D**). The effect was INSR-dependent, as S6 and ERK phosphorylation in response to insulin stimulation were attenuated by *INSR* knockdown (**Supplementary Figure 2**).

### *INSR* contributes to mammary tumourigenesis in BC model mice

To directly investigate the function of INSR in mammary tumorigenesis *in vivo*, we generated two genetically engineered mouse models with conditional mammary tissue-specific deletion of *INSR,* concomitant with expression of either oncogenic *HER2* (*MMTV-neu-IRES-Cre*, or *NIC*) or the polyoma middle-T (*PyMT*) antigen (*MMTV-rtTA;TetO-PyMT-IRES-Cre*, or *MTB:MIC*), respectively (**Figures 1E-F**) ^40–42^. In *NIC INSR* animals, *HER2* expression and *INSR* deletion are coordinated by the activity of the MMTV promoter, whereas in the *MTB:MIC INSR* model, MMTV promoter-dependent *PyMT* expression/*INSR* deletion are induced through treatment with doxycycline in the drinking water. Predicted allele and oncogene representation was monitored by genomic PCR (**Figures 1G-I**). Mammary tumours that developed in both models expressed INSR protein, while, consistent with their genotype, INSR was undetectable in mammary tumours of 10-week old *MTB:MIC INSR* and *NIC INSR* mice homozygous for the conditional *INSR* allele (F/F) compared to those wild-type for *INSR* (+/+) (**Figures 1J-K**). Metabolic INSR expressing tissues, such as liver, muscle, and fat, displayed unchanged INSR levels (**Figure 1K**), and maintained normal glucose homeostasis (**Figure 1L**), validating specificity of the mammary tissue *INSR* deficiency.

Following validation of oncogene expression and coordinate *INSR* deficiency in pilot transgenic experiments, cohorts were established of female mice carrying the respective oncogenes which were wild-type for *INSR*(+/+), *INSR* heterozygous-null (F/+), or *INSR* homozygous-null (F/F), and mice in both groups were monitored weekly for presence of tumours by palpation. In *NIC* mice, loss of *INSR* delayed tumour appearance in a deleted *INSR* allele-dependent manner, with *NIC INSR* (+/+) tumours emerging at 133 days, *NIC INSR* (F/+) at 158 days and *NIC INSR* (F/F) at 172 days (**Figure 2A**). Consistently, all *NIC INSR* (+/+) animals developed tumours within 172 days, while it took 294 days for all *NIC INSR* (F/F) mice to show signs of disease. In *MTB:MIC INSR* mice, where PyMT-Cre expression was induced at 10-12 weeks of age through doxycycline in the drinking water, PyMT induction rapidly led to appearance of tumours (∼8 days post induction) in all three *INSR* genotypes. While all *MTB:MIC INSR* (+/+) mice presented with tumours within 30 days post induction, at the 8-week monitoring endpoint, set due to the severity of the tumour phenotypes found in *MTB:MIC INSR* (+/+) mice, a considerable proportion of *MTB:MIC INSR* (F/F) and *MTB:MIC INSR* (F/+) mice either did not show signs of disease or had delayed tumour onset (**Figure 2B**). Reduction in tumours followed the number of deleted *INSR* copies, with >20% of *MTB:MIC INSR* (F/+) and >40% of *MTB:MIC INSR* (F/F) mice remaining tumour-free at endpoint.

**Figure 2:**
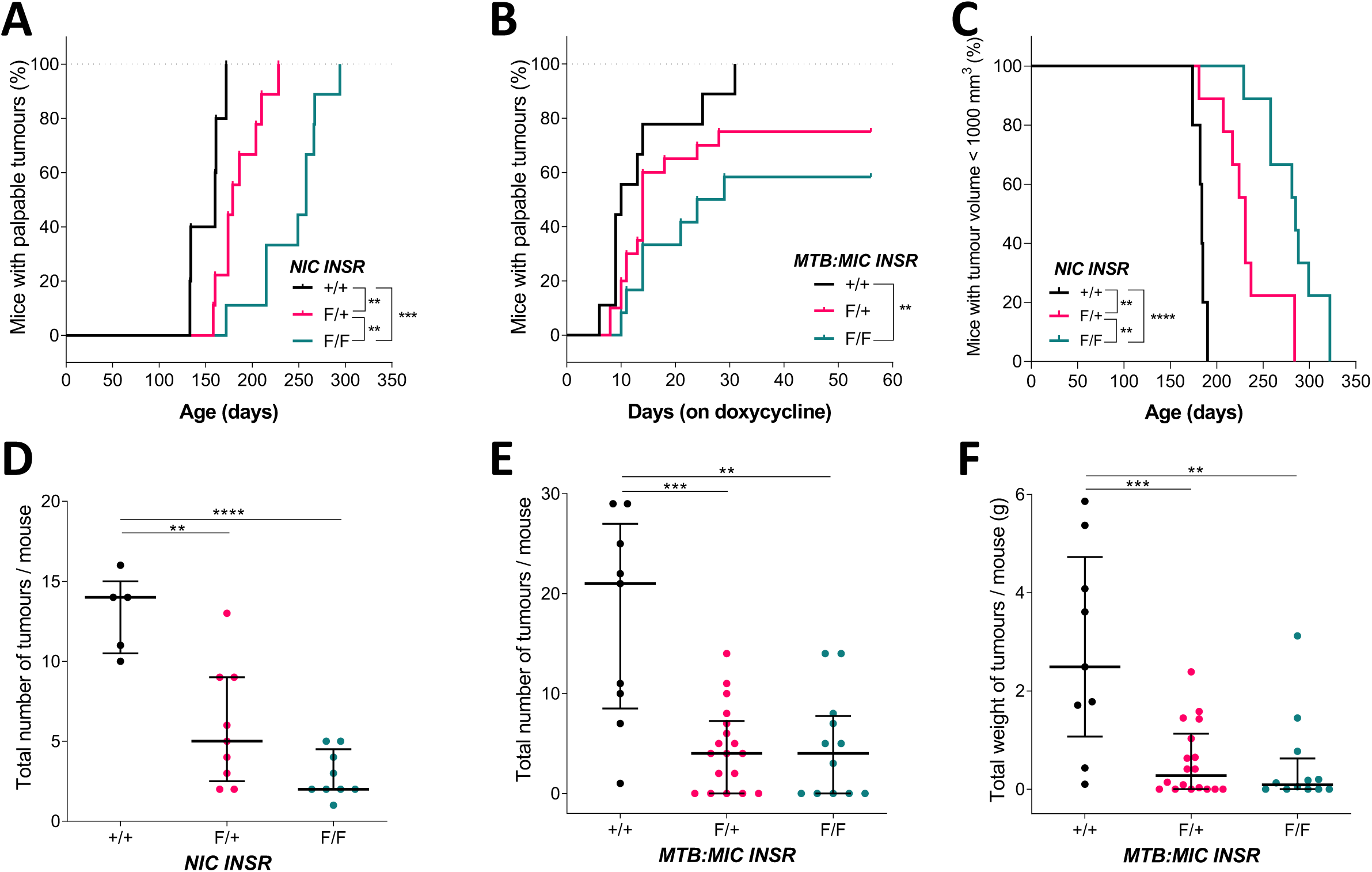
Loss of INSR delays mammary tumour onset, prolongs survival and reduces tumour burden in vivo. (**A**) Reverse Kaplan-Meier analysis of mammary tumour onset in *NIC* mice with wild-type *INSR* alleles (+/+, n = 5) and mice heterozygous (F/+, n = 9) and homozygous (F/F, n = 9) for the conditional *INSR* allele. (**B**) Reverse Kaplan-Meier analysis of mammary tumour onset (post-doxycycline regimen initiation) in *MTB:MIC* mice with wild-type *INSR* alleles (+/+, n = 9) and mice heterozygous (F/+, n = 20) and homozygous (F/F, n = 12) for the conditional *INSR* allele. (**C**) Kaplan-Meier analysis of overall survival (i.e. total tumour volume less than 1000 mm^3^) in *NIC* mice with wild-type *INSR* alleles (+/+, n = 5) and mice heterozygous (F/+, n = 9) and homozygous (F/F, n = 9) for the conditional *INSR* allele. (**D**) Scatterplot of endpoint tumour burden in *NIC INSR* mice (+/+, n = 5; F/+, n = 9; F/F, n = 9). Mice were sacrificed at humane endpoint (tumour volume exceeding 1000 mm^3^), followed by quantification of total tumour number. (**E-D**) Scatterplots of endpoint tumour burden in *MTB:MIC INSR* mice (+/+, n = 9; F/+, n = 18; F/F, n =12). Mice were sacrificed at 8 weeks post-doxycycline treatment regimen initiation, followed by quantification of total tumour (**E**) numbers and (**F**) weights. Scatterplots show the median and interquartile range. * p < 0.05, ** p < 0.01, *** p < 0.001, **** p < 0.0001; log-rank test (**A-C, I**), Student’s t-test (**D-F**).

Delayed tumour onset due to *INSR* deletion was also reflected in the time it took mice to reach the humane endpoint, as determined by overall tumour volume, with *NIC INSR* (F/+) and *NIC INSR* (F/F) mice displaying substantial survival benefit compared to their *NIC INSR* (+/+) counterparts (**Figure 2C**). Moreover, deletion of a single copy or both copies of *INSR* also led to remarkably lowered tumour burden in both models, reflected in the number and total weight of mammary tumours at endpoint. *NIC INSR* (+/+) mice developed up to 15 tumours per mouse, whereas *NIC INSR* (F/+) and *NIC INSR* (F/F) mice averaged around 5 lesions per mouse (**Figure 2D**). Similarly, *MTB:MIC INSR* (+/+) mice had >20 tumours per mouse on average, while both *MTB:MIC INSR* (F/+) and *MTB:MIC INSR (F/F)* mice featured ∼5 tumors per mouse at endpoint (**Figure 2E**). Reduced tumour count paralleled lower total tumour weights in *INSR-*deficient *MTB:MIC* mice at endpoint (**Figure 2F**).

### *INSR* deletion does not impact mammary tumour progression or metastasis

While *MTB:MIC INSR* tumours that escaped the impact of *INSR* loss appeared to, on average, weigh less at endpoint (**Figure 3A**), this reduction was not statistically significant and the extent of disease in this model precluded further comparisons of the impact on tumour progression. Nevertheless, slower tumour development in *NIC INSR* mice afforded a more detailed assessment of the impact of INSR on tumour progression. Diameters/volumes of individual tumors and total tumors per mouse in the three *NIC INSR* genotypes were recorded weekly by caliper measurements. Plotting of the growth of individual tumours and cumulative tumour volume per mouse over time suggested non-linear, exponential growth (**Figures 3B, E**). Data were log10 transformed (**Figures 3C, F**) and fit of the linear mixed models to both the original and log-transformed data were compared (see Methods). Comparison of percent growth per day values for each tumour calculated from the models (>10 per genotype) showed that there was no difference in progression of individual tumors between the genotypes (**Figure 3D**).

**Figure 3:**
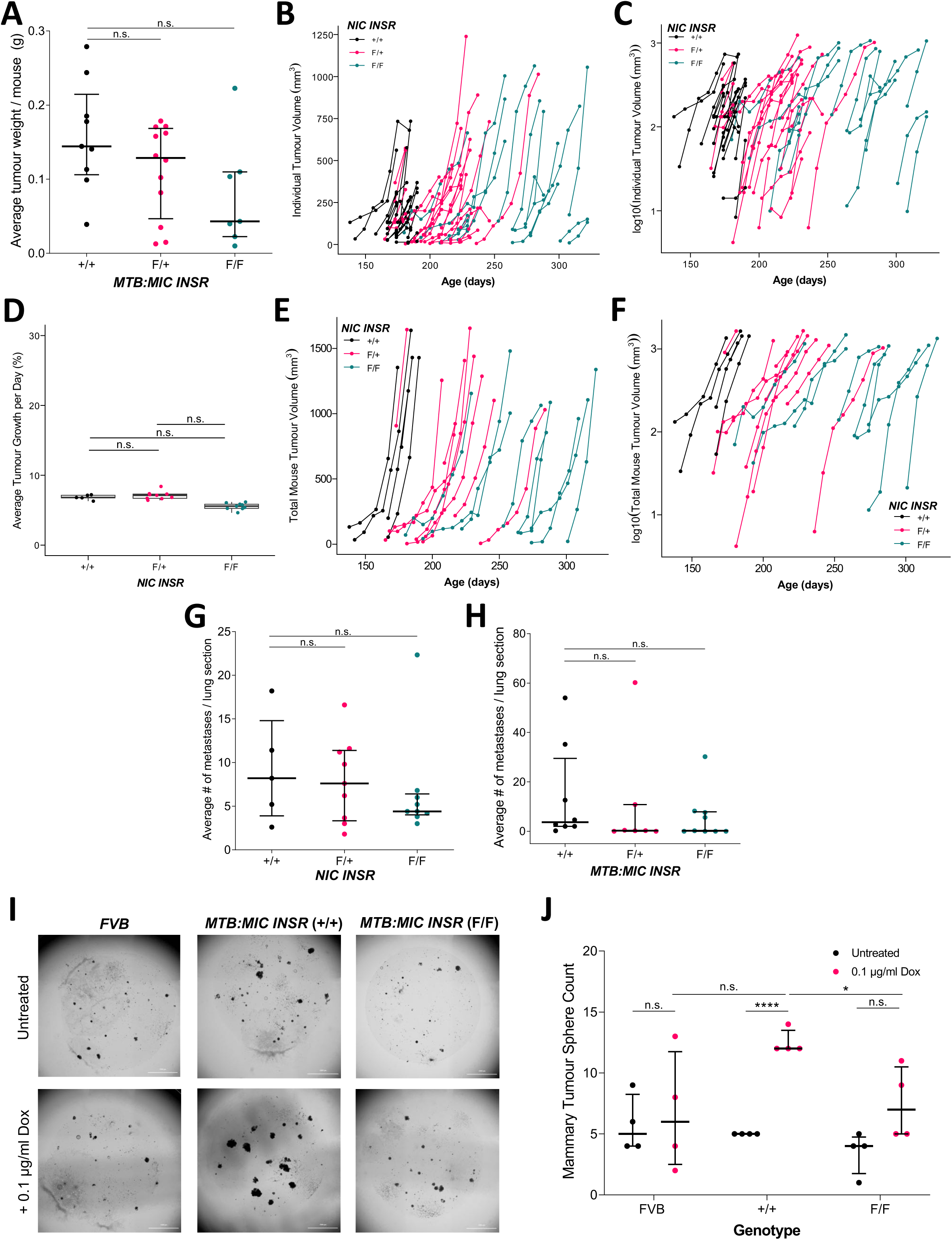
Loss of INSR impairs tumour initiation but does not impact tumour progression or metastasis. (**A**) Scatterplot of average weights of tumours at endpoint in *MTB:MIC INSR* mice which developed tumours by 8 weeks post-doxycycline induction. (**B-C**) Time-resolved plots for individual tumour growth in *NIC INSR* mice (+/+, n = 21; F/+, n = 24; F/F, n = 18) as a function of the age of the mouse in days, with tumour volume measurements displayed on (**B**) linear and (**C**) logarithmic scale. (**D**) Boxplot of estimated average % tumour growth per day for each *NIC INSR* mouse (+/+, n = 5; F/+, n = 9; F/F, n = 9). Estimates were calculated by adding together average growth rate for each genotype, and any estimated random variation from this average for each mouse. Values for each individual mouse are plotted over the boxplots. (**E-F**) Time-resolved plots for total tumour burden (as measured by summing total volume of each tumour per mouse) for *NIC INSR* mice (+/+, n = 5; F/+, n = 9; F/F, n = 9) as a function of the age of the mouse in days, with tumour volume measurements displayed on (**E**) linear and (**F**) logarithmic scale. (**G-H**) Scatterplots of metastatic burden in (**G**) *NIC INSR* mice (+/+, n = 5; F/+, n = 9; F/F, n = 9) and (**H**) *MTB:MIC INSR* mice RFD (+/+, n = 8; F/+, n = 7; F/F, n = 9). Mice were sacrificed at their respective endpoints and metastatic burden was assessed through quantification of the number of lung metastases as determined by H&E staining of FFPE lung sections, averaged across 5 sections. (I-**J**) Cells isolated from mammary glands of FVB, *MTB:MIC INSR* (+/+) *and MTB:MIC INSR* (F/F) mice were resuspended in Matrigel, cultured for 11 days in Epicult-B medium +/- 0.1 µg/mL doxycycline, followed by imaging using Cytation 5 live cell plate reader. (**I**) Representative Z-stack image projections of Matrigel domes. (**J**) Scatterplot of number of mammary tumour spheres per Matrigel dome as quantified using Gen 5 digital image analysis. Scatterplots show the median and interquartile range. Boxplots show the median and interquartile range, with Tukey style whiskers. * p < 0.05, **** p < 0.0001, n.s = non-significant; Student’s t-test (**A, G-H, J**), ANOVA (**D**).

To estimate the impact of *INSR* deletion on metastatic spread, number of lung metastases in both models were compared. In both the moderately metastatic *NIC INSR* and the highly metastatic *MIC:MTB INSR* models, deletion of either single or both copies of *INSR* had no effect on metastatic dissemination (**Figures 3G, H**).

### *INSR* loss impairs mammary tumour initiation

Despite *INSR*-deficiency delaying tumour onset and reducing endpoint tumour burden, the lack of effect of *INSR* loss on progression and metastasis points to the possibility that the functional contribution of INSR to BC occurs at an earlier time point in tumorigenesis. To directly probe the impact of *INSR* deletion on tumor initiation, mouse mammary epithelial cells from *MIC:MTB INSR* (+/+) and *MIC:MTB INSR* (F/F) not treated with Dox, and thus equivalent, were isolated and grown in Matrigel (**Figure 3I**). Tumour sphere formation was induced by continuous treatment with Dox (0.1 µg/mL) for 11 days and the appearance of tumour spheres were quantified. PyMT induction led to abundant tumor sphere formation by *MIC:MTB INSR* (+/+) mammary cells compared to those from parental *FVB* mice, or Dox-untreated controls (**Figure 3J**). By contrast, *INSR* deletion led to significantly lower tumor sphere development, supporting the notion that *INSR* loss reduces tumour initiation.

### *INSR*-deficient tumours maintain canonical pathway signaling activity

To gain better insight into the lack of impact of INSR signaling on progressing mammary tumours *in vivo*, immunoblotting of tumor lysates and immunohistochemistry of tumor tissue sections collected at endpoint were performed in both models (**Figure 4; Supplementary Figures 3-6**). Immunoblotting for signaling pathway activity in tumour lysates failed to yield consistent output (**Supplementary Figure 3**), thought to be affected by the presence of stromal and other non-tumour tissue content in tumour isolates.

**Figure 4:**
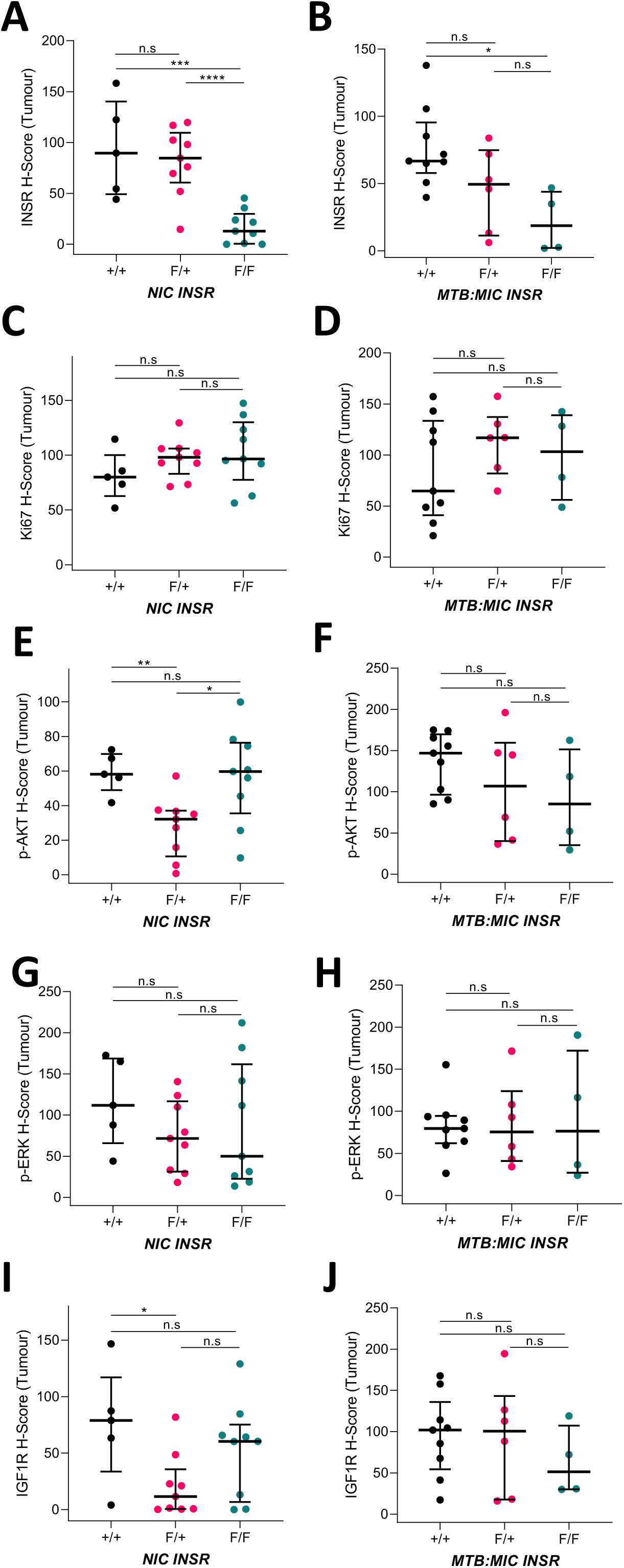
INSR-deficient tumours display no difference in canonical signaling downstream of INSR. Scatterplots of digital pathological analysis using QuPath software of immunohistochemically stained sections of paraffin embedded tumours collected at endpoint from (**A**, **C**, **E**, **G**, **I**) *NIC INSR* mice (+/+, n = 5; F/+, n = 9; F/F, n = 9), (**B**, **D**, **F**, **H**, **J**) *MTB:MIC INSR* mice fed RFD (+/+, n = 9; F/+, n = 6; F/F, n = 4). H-scores for scanned images of stained tumour sections were determined by QuPath using *Simple Tissue Detection* followed by *Positive Cell Detection* and tissue classification, which selectively scored individual tumour cells as positive (1+, 2+ or 3+) or negative (0) for staining of (**A-B**) INSR, (**C-D**) Ki67, (**E-F**) p-AKT, (**G-H**) p-ERK, and (**I-J**) IGF1R. Scatterplots show the median and interquartile range. * p < 0.05, ** p < 0.01, *** p < 0.001, **** p < 0.0001, n.s. = not significant; Student’s t-test.

To quantitatively assess the tumour-specific biology of collected tissues, immunohistochemically stained sections of endpoint tumours were digitized (**Supplementary Figure 4**) and analyzed (**Figure 4; Supplementary Figures 5-6**). In agreement with their genotype, QuPath analysis^43^ revealed that tumour-specific cells of *NIC INSR* (F/F) and *MTB:MIC INSR* (F/F) mice had greatly diminished INSR staining relative to their *INSR* (+/+) counterparts, with the tumour cells of mice heterozygous for *INSR* loss (F/+) exhibiting intermediary but non-statistically significant reduction (**Figures 4A-C; Supplementary Figures 5A-C**). Consistent with no impact on tumour progression, Ki67 staining, a marker of active proliferation in tumour cells, remained comparable at endpoint across *INSR* genotypes of both models (**Figures 4D-F; Supplementary Figures 5D-F**). This was paralleled by analogous lack of significant change in levels of phosphorylated AKT (**Figures 4G-I; Supplementary Figures 5G-I**) and phosphorylated ERK (**Figures 4J-L; Supplementary Figures 5J-L**). Potential compensatory changes in IGF1R levels in response to genetic *INSR* disruption were also tested by IHC to reveal no difference between genotypes (**Figures 4M-O; Supplementary Figures 5M-O**). Of note, *NIC INSR* (F/+) tumours displayed statistically significant reduction of both p-AKT (**Figure 4G; Supplementary Figure 5G**) and IGF1R (**Figure 4M; Supplementary Figure 5M**), an observation that was not accompanied by decreased proliferative signal (**Figure 4D; Supplementary Figure 5D**), nor with further reduction tracking with INSR levels in *NIC INSR* (F/F) tumours (**Figure 4A; Supplementary Figure 5A**). Judging by immunohistochemistry for the pan-immune marker CD45, there was no difference between *INSR* genotypes in the general immune infiltrates of *MTB:MIC INSR* tumours collected at 10 weeks post-doxycycline induction (**Supplementary Figure 6**).

### *INSR* loss in murine mammary epithelial cells impairs their bioenergetic potential

To gain further insight into the mechanism(s) behind reduced tumour initiation upon *INSR* deletion, we generated two independent *INSR*-deficient luminal COMMA1D mouse mammary cell lines (COMMA1D INSR (-/-)) and two accompanying wild-type controls (COMMA1D INSR (+/+)) using CRISPR/Cas9 (**Supplementary Figure 7A**) ^44^. COMMA1D cells are immortalized but not transformed mammary epithelial cells isolated from *Balb/C* mice in mid-pregnancy and have been used to study oncogenic mammary transformation ^45,46^. *INSR* deletion in COMMA1D cells did not affect net activation of the PI3K-Akt (**Supplementary Figures 7B, C**) and Ras-MAPK (**Supplementary Figures 7B, D**) signaling pathway components in response to insulin stimulation, likely due to the presence of IGF1R, as evidenced by a reduced but not depleted phospho-INSR/IGF1R signal in *INSR*-deficient cells (**Supplementary Figure 7B, E**).

COMMA1D INSR cells were further characterized using RNAseq. Based on GSEA, mRNA expression of genes associated with DNA replication, cell cycle regulation and mRNA translation/protein synthesis were affected by *INSR* deletion (**Supplementary Figure 8A**). Notably, we also observed INSR-dependent differential expression of a number of genes involved in mitochondrial respiratory electron transport (**Figure 5A; Supplementary Figures 8B-D**). More specifically, expression of many genes encoding components of respiratory complexes I-IV and complex V (ATP synthase) were lowered in *INSR*-deficient cells compared to their wild-type counterparts (**Supplementary Figure 8D**). Differential mRNA expression was corroborated by RT-qPCR of the select components of each respiratory complex (**Figure 5B**). To further validate these observations, equivalent analysis was performed in the independent GSE206565 RNAseq dataset ^47^, in which mouse preadipocytes lacking both *INSR* and *IGF1R* (double knock-out, DKO) were reconstituted with either wild-type INSR (IR), a kinase-dead INSR mutant (KD), a truncated INSR lacking the C-terminus (CT), or a truncated INSR lacking the majority of the intracellular domain (juxtamembrane-domain only, JMO). Similar to the observations in our COMMA1D cells, dual *INSR*/*IGF1R* loss in these cells resulted in reduced expression of mitochondrial respiratory chain complex components, particularly in complexes I and IV (**Supplementary Figure 9**). Markedly, downregulation of subunits of these respiratory chain complex components in DKO cells was rescued by reconstitution of the kinase-dead INSR mutant, implicating non-canonical/ligand-and-tyrosine-kinase-independent (LYK-I) INSR signaling in mediating this effect.

**Figure 5:**
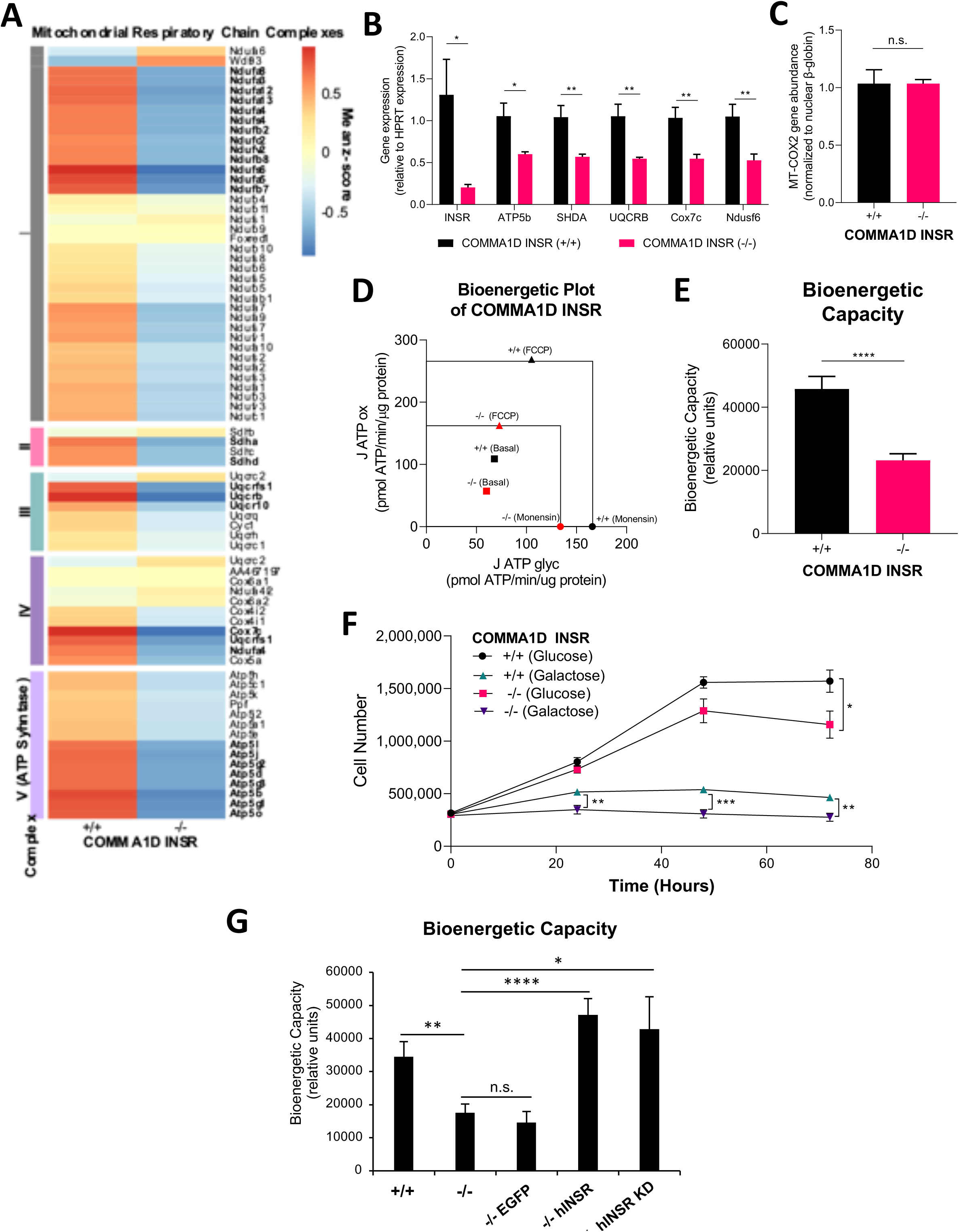
INSR loss in mammary epithelial cells reduces bioenergetic capacity in kinase activity-independent manner. (**A**) Heatmap of mean z-score normalized expression by RNAseq for replicates of COMMA1D INSR (+/+) cells (n = 6) and COMMA1D INSR (-/-) cells (n = 6) for genes associated with mitochondrial respiratory chain complexes. Genes differentially expressed (FDR < 0.05) between genotypes are highlighted in bold. (**B**) Bar graph showing relative mRNA expression by RT-qPCR of *INSR* and select components of the mitochondrial respiratory chain shown to be differentially expressed in COMMA1D INSR cells in **A**. Gene expression levels were normalized to *HPRT* expression. (**C**) Bar graph showing the relative mitochondrial DNA content in COMMA1D cells, as measured by qPCR of MT-genome encoded *COX2* (mitochondrial gene) normalized to nuclear genome-encoded β-globin. (**D-E**) Seahorse Mito Stress Test Assay of COMMA1D INSR (+/+) (n = 39) and COMMA1D INSR (-/-) (n = 40) cells seeded onto 96-well Seahorse assay plates. Compounds were loaded into a hydrated cartridge in the following order: Port A 1 µM Oligomycin, Port B 2µM FCCP, Port C 1 µM Rotenone/ Antimycin A, and Port D 20 µM Monensin. Measurements were determined by WAVE software. (**D**) Bioenergetic plot representing the average glycolytic ATP production rate (J ATP-glyc; x-axis) plotted against the average mitochondrial respiratory ATP production rate (J ATP-ox; y-axis) of COMMA1D INSR cells under basal, maximal OXPHOS (FCCP), and maximal glycolytic (Monensin) conditions. Areas under the boxes represent average bioenergetic capacity. (**E**) Bar graph representing bioenergetic capacities from **D**. (**F**) Glucose/Galactose assay of COMMA1D INSR (+/+) (n = 6) and COMMA1D INSR (-/-) (n = 6) cells seeded into 6-well plates with DMEM/F12 glucose-free media supplemented with glucose and counted the following day, as well as counted the next three days following media replenishment using DMEM/F12 glucose-free media supplemented with either 17 mM glucose (permits either oxidative phosphorylation or glycolysis) or 17 mM galactose (permits oxidative phosphorylation only). (**G**) COMMA1D INSR (+/+) and (-/-), and COMMA1D INSR (-/-) expressing EGFP, wt INSR or KD INSR K1030R were analyzed as in D-E. Bar graphs show mean with error bars representing SEM. Time-resolved plot shows mean with error bars representing SEM. * p < 0.05, ** p < 0.01, *** p < 0.001, **** p < 0.0001, n.s. = non-significant; Student’s t-test.

We tested whether the deletion of *INSR* in COMMA1D cells resulted in alterations in cellular mitochondrial content. qPCR-DNA analysis of mitochondrial genome-encoded *COX2* gene levels relative to that of nuclear encoded β-globin indicated no differences in mitochondrial content between the *INSR*-deficient and - proficient cells (**Figure 5C**). We next investigated whether *INSR* deletion-dependent changes in mRNA abundance of respiratory chain components affect cellular bioenergetics by performing respirometry analysis (**Figures 5D; Supplementary Figure 10**). Monitoring of oxygen consumption rate (OCR) revealed diminished mitochondrial respiration in COMMA1D INSR (-/-) cells relative to their wild-type counterparts (**Supplementary Figure 10A**), evidenced by reduced basal and maximal respiration rates and ATP generated by oxidative phosphorylation (J-ATP ox) (**Figure 5D; Supplementary Figure 10C-D**). Parallel measurements of extracellular acidification rates (ECAR) (**Supplementary Figure 10B**) revealed reduced maximal glycolysis and ATP generated by glycolysis (J-ATP glyc) in *INSR* knockout cells compared to controls (**Figure 5D; Supplementary Figure 10C, E**). Together, *INSR* loss led to a major reduction of total bioenergetic capacity of COMMA1D cells (**Figure 5D, E**). In concert with restoration of mRNA expression of mitochondrial respiratory chain components in the preadipocyte model by both the WT the INSR kinase-dead mutant (**Supplementary Figure 9**), unlike control EGFP, re-expression of the wt INSR and the kinase-dead INSR K1030R mutant rescued the bioenergetic phenotype of INSR-deficient Comma1D cells (**Figure 5G**, **Supplementary Figure 10 F-G**).

To further explore the biological significance of the bioenergetic impairment of *INSR*-deficient cells, glucose was replaced by galactose as the carbohydrate source in the growth media, forcing cells to rely on mitochondrial oxidative phosphorylation for ATP production ^48,49^. Monitored over 3 days, COMMA1D INSR (-/-) cells failed to proliferate in galactose-containing media (**Figure 5F**), further corroborating gene expression (**Figure 5A; Supplementary Figure 8**) and respirometry findings (**Figure 5D, E; Supplementary Figure 10**).

## Discussion

Human epidemiological and observational studies have defined a strong correlation between excess body weight and incidence and burden of many cancers, including BC. Obesity is a key driver in cardiovascular, musculoskeletal and kidney diseases, as well as a causal factor in the development of T2D ^50^. The rapidly rising global prevalence of obesity makes it one of the greatest threats to human health in the 21^st^ century. There are a number of systemic and molecular mechanisms implicated in mediating the harmful effects of obesity on human health, including the development of cancers ^51^. Coupled with broad expression of INSR in human cancers (**Figure 1A**), hyperinsulinemia, which correlates with worse BC outcomes ^7,52,53^, is one of the strongest obesity corollaries thought to contribute to cancer burden (reviewed in ^54^). The relationship between obesity, hyperinsulinemia, T2D, menopausal status and BC is complex ^55,56^. While increased adiposity in postmenopausal women associates with an increase in BC risk, obesity and T2DM appear to have either no effect or even some protective effect in premenopausal women ^57–60^.

Using two well-defined genetically engineered mouse models of BC driven by the *PyMT* and *HER2* oncogenes, respectively, we directly probed the importance of the INSR to the development of mammary tumours by its coordinate genetic deletion in tumour tissue. Transcriptional profiling classifies PyMT mammary tumors with the human BC luminal B subtype ^61^, whereas mammary tumours in the HER2 transgene model recapitulate many of the features of HER2-positive human BCs ^62^. Control tumours from both models express INSR (**Figures 1J-K**), a feature found in >80% of human BC ^18,19^.

In both models, deletion of a single or both *INSR* copies dramatically attenuated tumor onset, leading to reduced tumor burden at endpoint (**Figures 2A-F**). Longitudinal monitoring of individual tumours in the HER2 model revealed that once tumours appeared, *INSR* deficiency had no impact on progression or metastasis (**Figures 3A-H**), indicating that INSR likely participates in the tumour initiation phases. Indeed, when tumor spheres were initiated by PyMT *ex vivo*, *INSR* deletion significantly impaired their formation (**Figures 3I-J**), further corroborating the importance of INSR at the earliest stages of mammary tumorigenesis.

Mechanistically, mammary tumours driven by PyMT result from association-dependent activation of RTK-proximal effectors of mitogenic PI3K-Akt and Ras-MAPK signaling pathways ^63,64^ and the same pathways are also activated by superphysiological transgenesis of HER2 ^62^. From the tumour analysis at endpoint, *INSR* loss did not affect measures of proliferation, nor the PI3K-Akt and Ras-MAPK signaling throughput, highly representative of a tumour’s stage of progression rather than *INSR* deficiency (**Figure 4; Supplementary Figure 5**). Complementary studies explored the effects of hyperinsulinemia without obesity on PyMT-driven mouse xenograft growth and found increased insulin-dependent acceleration of the development and progression of tumours ^15^, whereas the same conditions augmented tumor development and metastasis of HER2-driven xenografts ^65^, accompanied by hyperactivation of Akt signaling in both models. Moreover, shRNA-mediated interference with *INSR* in BC cells in culture led to reduced proliferation and Akt signaling ^66,67^. Thus, while hyperinsulinemia can accelerate mammary tumour progression and metastasis in the context of PyMT and HER2 transgenesis in xenografts and *INSR* deletion can reduce *in vitro* BC cell line growth, presumed diminution of insulin signaling via reduction of INSR levels in mouse mammary gland tumours does not translate to effects on proliferation nor PI3K-Akt/Ras-MAPK pathway control at later stages of tumorigenesis. The differences between models might be a reflection of a developed insulin/INSR dependence of cultured and xenografted cells while grown *in vitro* and subcutaneously, or distinct tumor heterogeneity in the absence or reduced INSR since inception in the mouse genetically engineered models.

There is evidence that IGF1R and INSR can compensate for each other under conditions of pharmacological intervention or deficiency ^66,68^, as well as during mouse embryonic development ^69^. In our *MTB:MIC INSR* mouse models, levels of IGF1R remained unchanged following *INSR* loss (**Figures 5N-O; Supplementary Figures 5N- O**), suggesting lack of possible compensatory function of IGF1R in maintaining tumour proliferation and Erk/Akt signaling when *INSR* is reduced/deleted in PyMT-driven models. However, in *NIC INSR* mice, heterozygous *INSR* loss resulted in significantly reduced IGF1R expression (**Figure 4M; Supplementary Figure 5M**) and Akt signaling (**Figure 4G; Supplementary Figure 5G**), an effect not observed with homozygous *INSR* loss. Given that INSR and IGF1R can also form hybrid receptors ^30^, AKT signaling in endpoint HER2-driven tumours may be dependent on INSR under normal conditions, but this dependency shifts to hybrid receptors or IGF1R holo-receptors under conditions of reduced INSR or complete INSR deficiency, respectively. Regardless, high proliferation rates and Erk/Akt activity-associated phosphorylation upon *INSR* loss were paralleled by the unchanged progression indicators in *INSR*-deficient tumours, with both likely originating from the *PyMT* and *HER2* transgenes. Thus, *INSR* deletion impacts the early PyMT- and HER2-driven steps towards mammary epithelial transformation and initiation of primary tumours, but not the later phases of progression and metastasis.

Bioenergetic plasticity of cancer cells has long been recognized as one of neoplastic hallmarks (reviewed in ^70^). While proliferating cancer cell’s carbon and energy hunger that accompany new biomass production, and reliance on aerobic glycolysis for energy production (the Warburg effect) ^71,72^ have been well studied, less is known about the energetic landscape of pre-cancerous cells during tumour initiation. Our mRNA expression analyses in *INSR*-deficient non-transformed mammary epithelial cells indicates that *INSR* loss leads to reduced expression of many subunits of the mitochondrial respiratory chain (MCR) complexes (**Figure 5A; Supplementary Figure 8**), an observation validated by analysis of the data from a complementary study in mouse preadipocytes (**Supplementary Figure 9**). Further analyses of the INSR- deficient COMMA1D cells revealed *INSR* loss-dependent breakdown in bioenergetic capacity (**Figure 5E**), through diminished ATP production via both OXPHOS and glycolysis (**Figures 5C, E**). Indeed, when challenged to grow in galactose as a carbohydrate source, which necessitates the use of OXPHOS for energy production, *INSR*-deficient cells failed to thrive (**Figure 5J**). The same cells upregulated mRNA expression of genes belonging to GO terms defining carbohydrate starvation and transport, as well as amino acid starvation, consistent with their bioenergetic deficiency (**Supplementary Figures 8B-C**). Remarkably, parallel to the reconstitution of impaired MCR complex mRNA expression seen in mouse preadipocytes lacking both *INSR* and *IGF1R* by re-expression of a kinase-dead INSR mutant (**Supplementary Figure 9**), bioenergetic capacity of INSR-deficient Comma1D cells was rescued by both WT and kinase-dead INSR (**Figure 5G; Supplementary Figure 10 F, G**). Taken together, these findings raise the possibility of a generalizable function of ligand-and-tyrosine-kinase independent (i.e., non-canonical) INSR signaling mechanism(s), to uphold cellular bioenergetic fitness and enable neoplastic transformation.

Several lines of evidence suggest that insulin/INSR contribution to tumorigenesis might be particularly prominent in cells/tissues sensitized to insulin action by acquisition of mutations impacting the PI3K-Akt and Ras-MAPK signaling pathways (reviewed in ^25^). In our model systems, interference with INSR did not alter progression or metastasis of tumours caused by major drivers of these pathways, PyMT and HER2. Nevertheless, our work has revealed a parallel, likely kinase activity-independent mechanism of INSR- mediated tumour initiation. The comparable reduction in tumour burden by deletion of a single or both copies of *INSR* in our models indicates that there is a threshold for the contribution of INSR to tumour initiation. Further work aimed at reconciling the distinct mechanistic paths of INSR function in tumorigenesis will inform prevention and therapeutic strategies aimed at cancers featuring INSR expression and/or hyperinsulinemia.

### Limitations of Study

Our work reveals a previously unrecognized INSR function in mammary tumour initiation and tumorigenesis but presents with certain limitations. Tumor initiation is one of the least explored aspects of tumorigenesis. Paucity of what can be defined as cancer initiating cell(s) limits the experimental approaches that can be applied towards understanding of the molecular mechanisms, an area that can be advanced through precision genetics and single cell analysis of early lesions. We employed two mammary tumour models that recapitulate features of the majority of human BC, but not the triple-negative subtype (10-15% of all BC). Existing triple-negative BC models require conditional deletion of 2 tumour suppressor genes, often BRCA1 and p53 ^73^, which combined with coordinate tissue specific Cre-recombinase expression and INSR deletion, renders the desired genotype in female mice scarce, and any subsequent study currently prohibitively resource- and time-intensive.

While previous observational and animal/cell model work firmly considered insulin/INSR as an “axis”, we found that INSR might be impacting early tumorigenesis acting independently of its kinase activity and acting via mitochondria. Dependence of the observed effects on insulin stimulation, mechanism(s) of INSR bioenergetic control and factors downstream and beyond mitochondria warrant further study, in animal models *in vivo*, and through retrospective and prospective observational studies of early human breast metaplastic and neoplastic lesions, including ductal carcinoma *in situ*, from normal and hyperinsulinemic individuals. The existence of an INSR threshold for contribution to cancer also entails additional probing of the relevance of the insulin/INSR relationship in tumorigenesis, via genetic manipulation of epithelial INSR levels in healthy and tumour models, with parallel reduction or increase in systemic insulin levels ^16^, as well as under conditions mimicking the effects of obesity. Moreover, precision genetic animal model approaches aimed at uncoupling kinase activity-dependent and - independent epithelial cell INSR functions will further advance the underlying mechanistic understanding. Finally, recognition of the newly discovered INSR-upheld bioenergetic fitness in mammary epithelial cells, previously seen in metabolic insulin-target cells and tissues ^74^, as a potential rate limiting feature of neoplastic transformation, requires consideration when designing lifestyle and therapeutic interventions aimed at improving metabolic health as a cancer prevention measure.

## Acknowledgements

We would like to thank the staff in the Animal Research Center at the University Health Network (UHN) for help with animal husbandry, UHN’s Applied Molecular Profiling Laboratory for assistance with immunohistochemistry and Soumili Sarkar for help with image acquisition. We are indebted to Drs. David Papadopoli and Ivan Topisirovic for their help with bioenergetic analyses. This work was supported by the Canadian Institutes of Health Research Project Grant (162296) and the Canadian Cancer Society Grant (i2I16-2 grant #704786) to V.S. L.P. is a recipient of The Princess Margaret Hospital Foundation Graduate Fellowships in Cancer Research. Y. P. was a recipient of the Canadian BC Foundation postdoctoral fellowship and R. D. of the Banting Postdoctoral Fellowship from the Government of Canada.

## Author Contributions

Experiments were designed by Y.P., L.P. and V.S. and performed by Y.P., L.P., and C.I. S.M and S.L. helped with animal husbandry. Data, gene expression and statistical analyses was performed by L.P, M.E., P.S., and K.N. B.Z. S.S. and B.M. participated in data acquisition. R.D, F.V. and P.T. provided conceptual and technical input. L.P., Y.P. and V.S. wrote the manuscript. All authors discussed the results and commented on the manuscript.

## Inclusion and Diversity

We support inclusive, diverse, and equitable conduct of research.

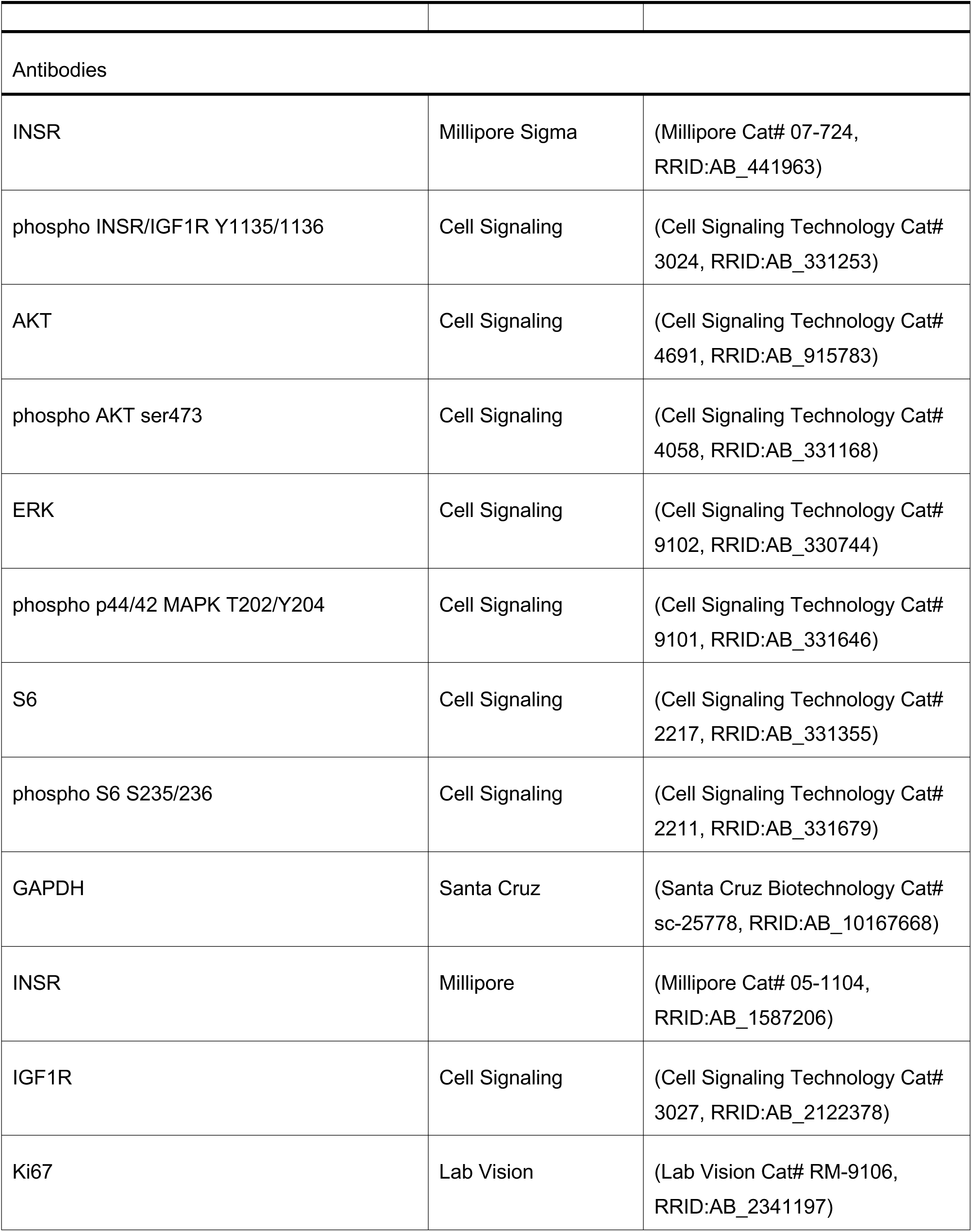
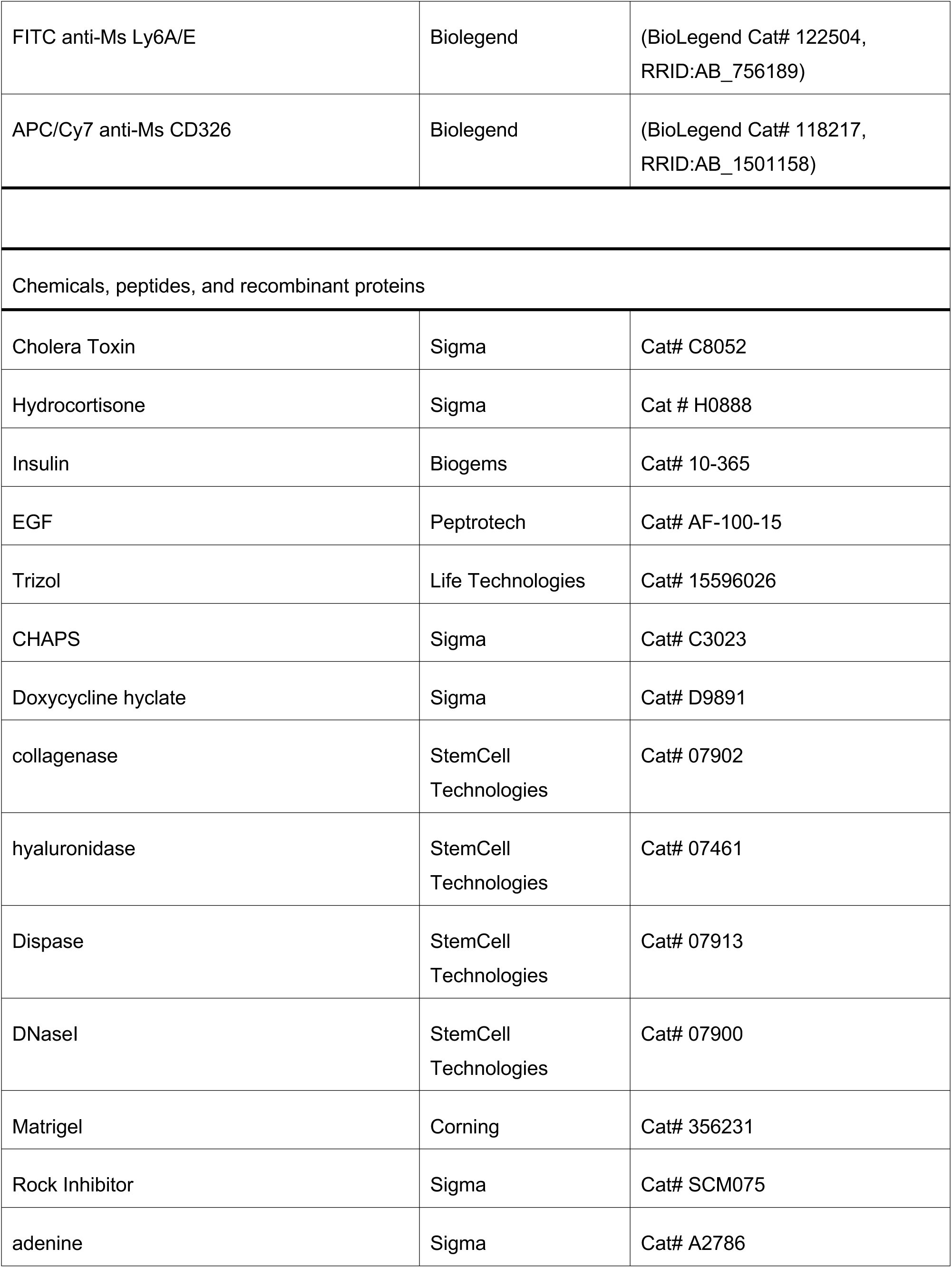
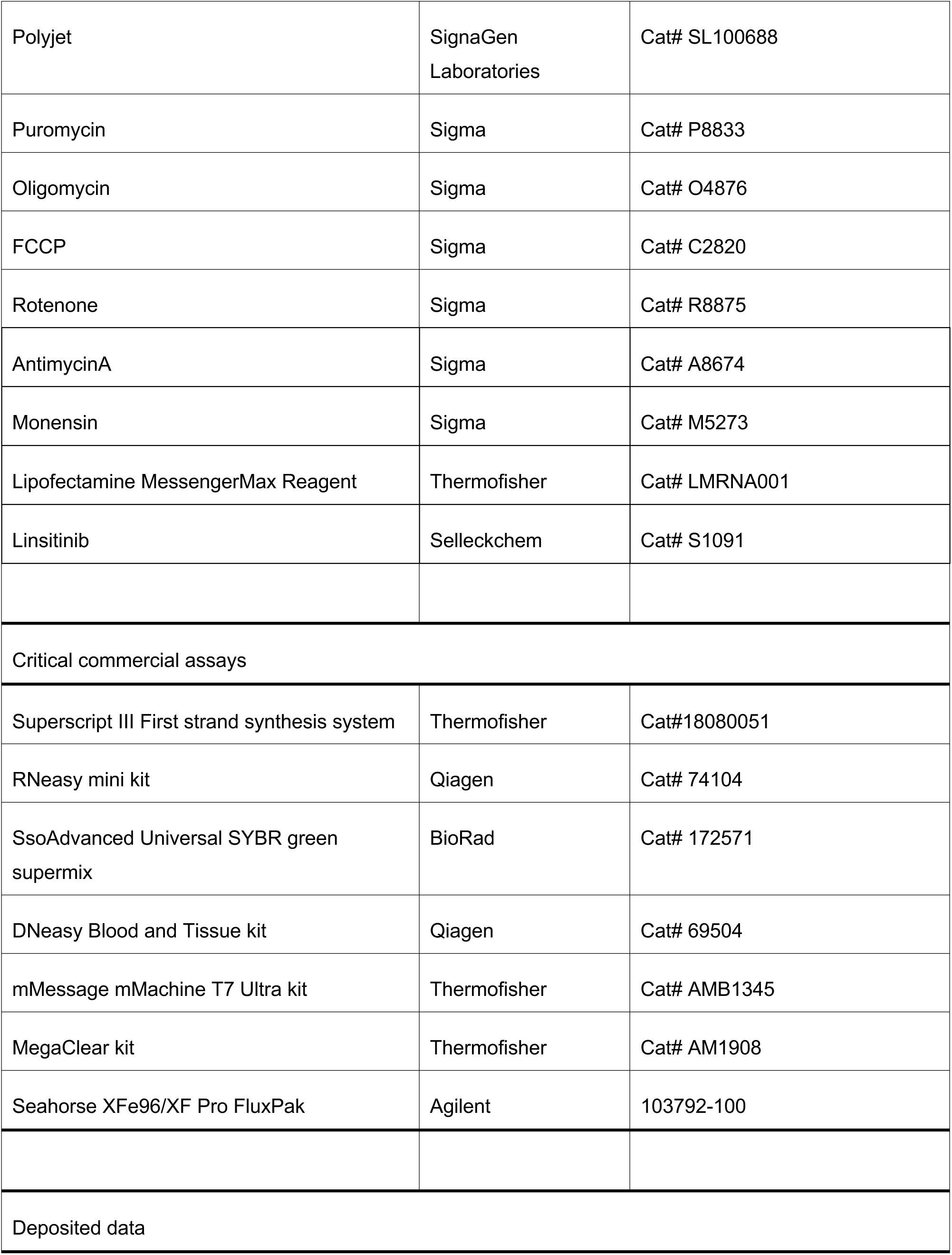
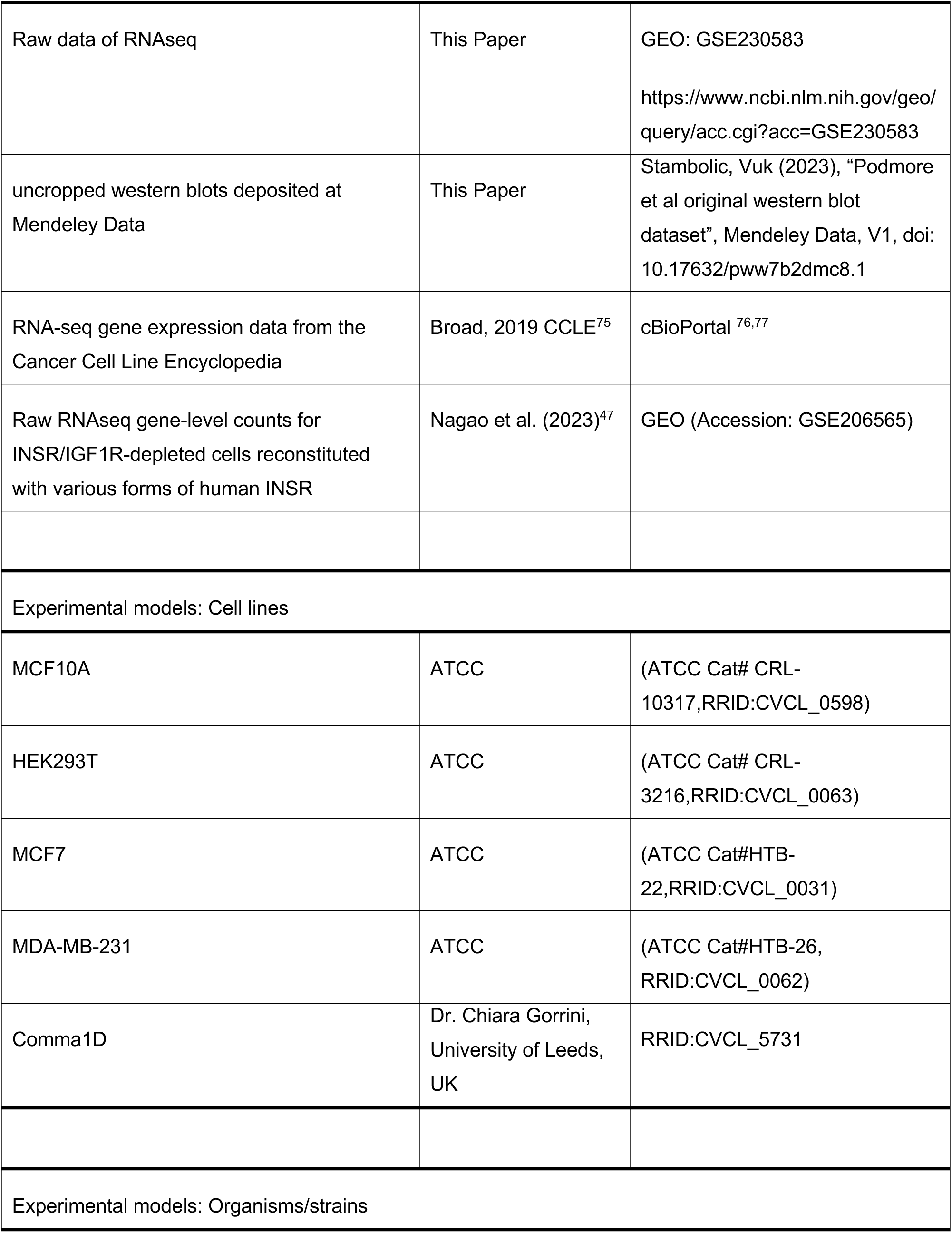
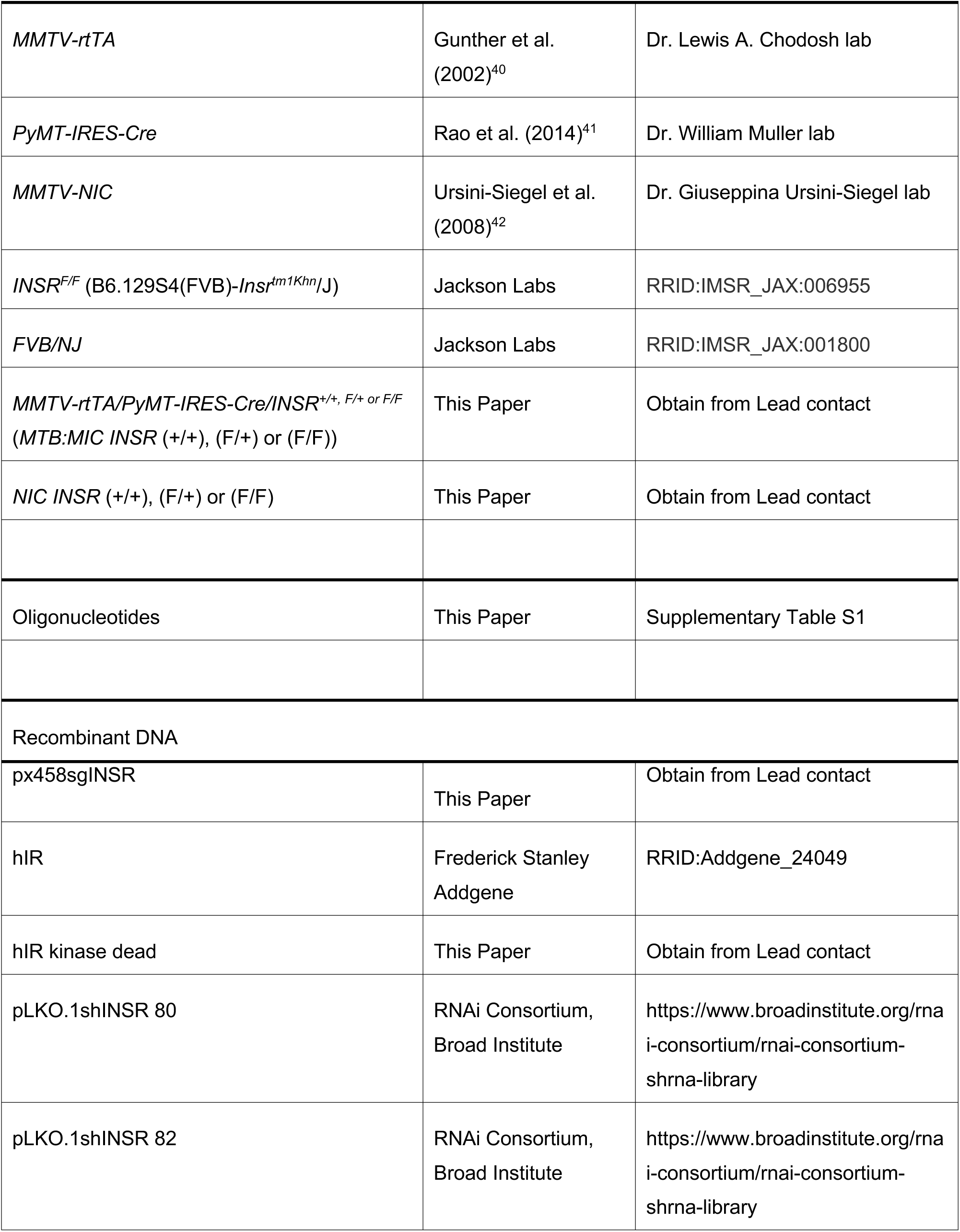
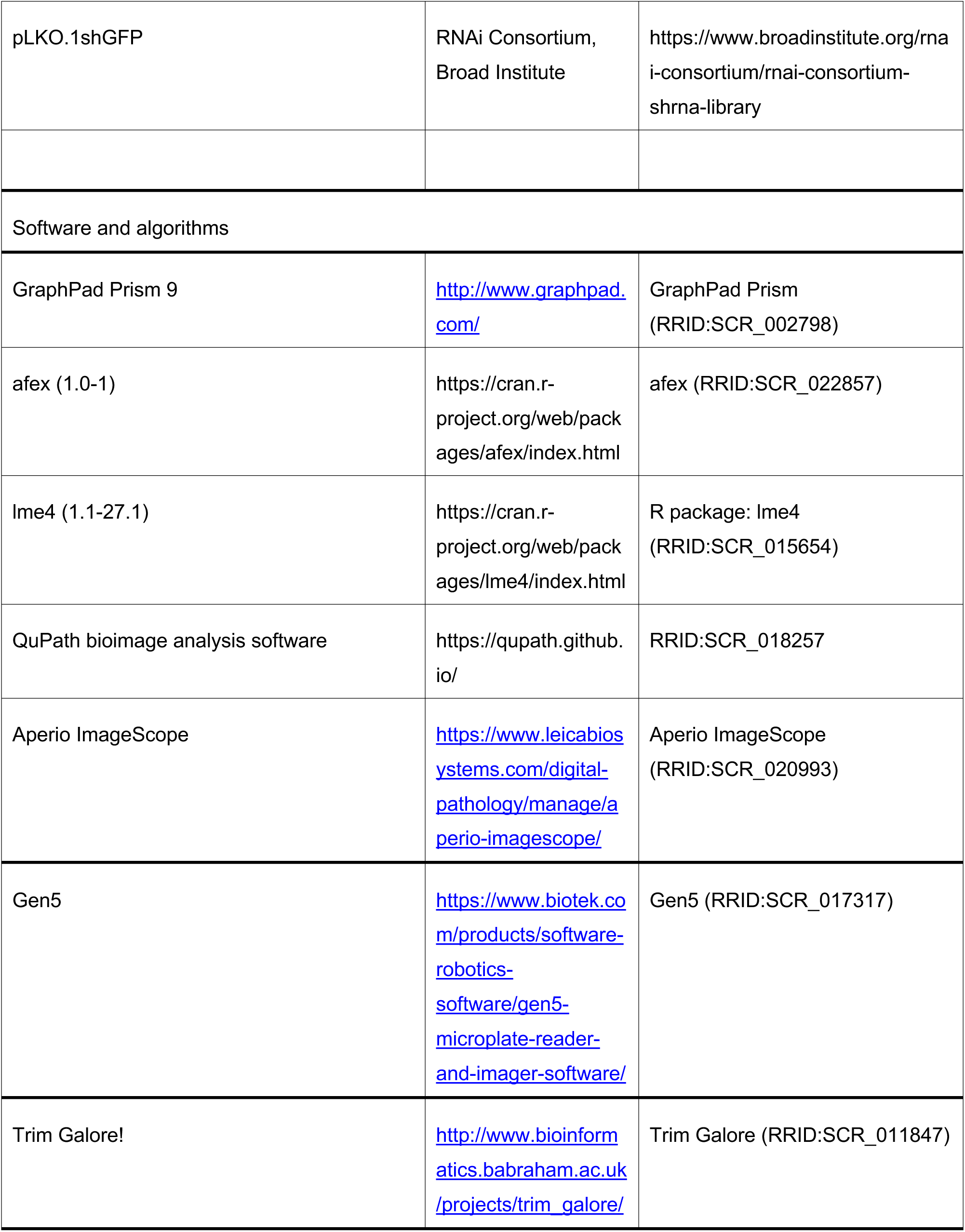
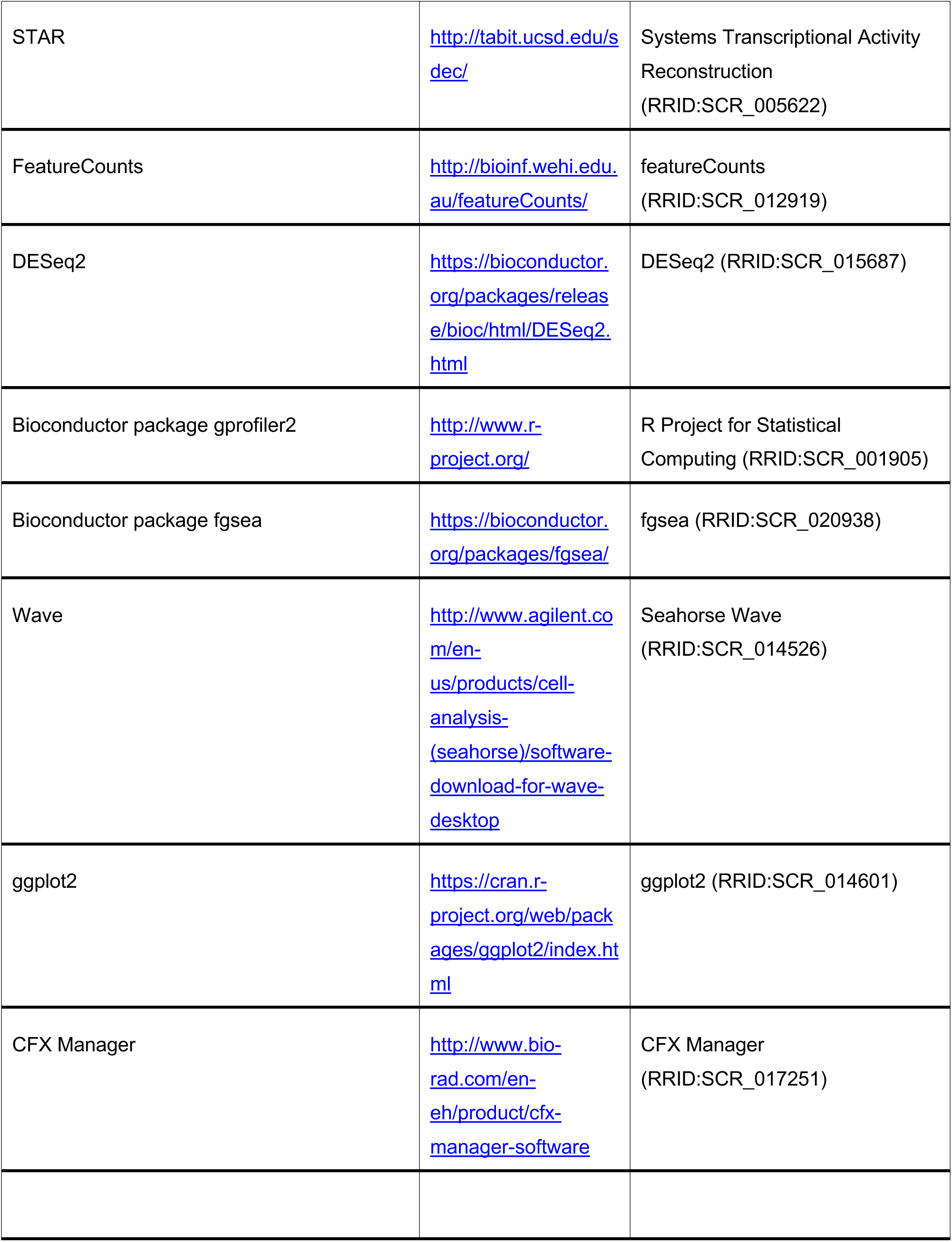
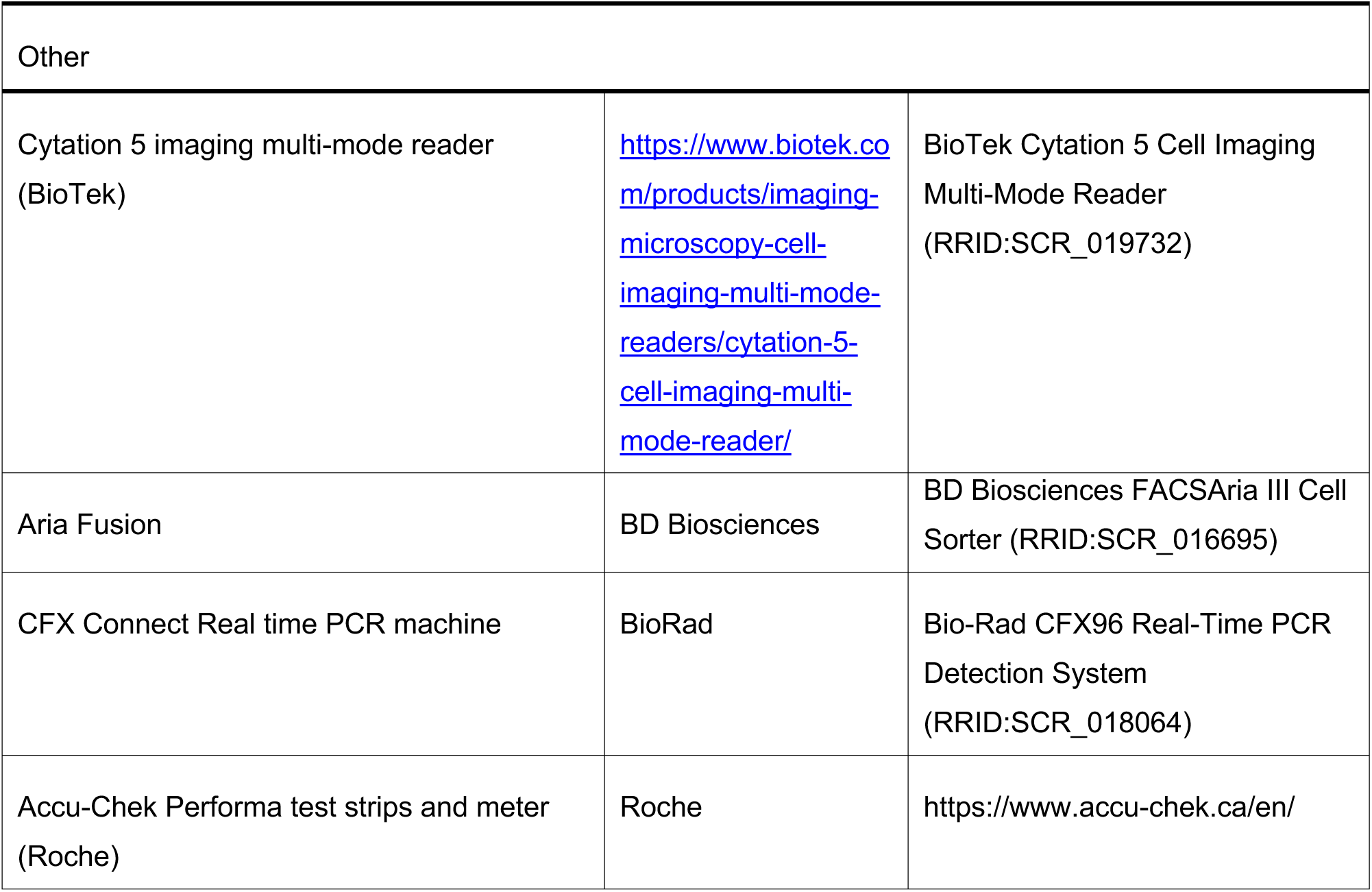
Key Resources Table.

## Resource availability

### Lead contact

Further information and requests for resources and reagents should be directed to and will be fulfilled by the lead contact, Vuk Stambolic.

### Materials availability

- Plasmids generated in this study are available from the lead author.
- INSR^F/F^ (B6.129S4(FVB)-Insr^tm1Khn^/J; stock #006955) were backcrossed onto the FVB/NJ mice (stock #001800) for 6 generations to generate INSR^F/F^ FVB/NJ mice. These mice are available from the lead author’s laboratory.

### Data and code availability

- RNA-seq data have been deposited at GEO (GEO GSE230583) and are publicly available as of the date of publication. Original western blot images have been deposited at Mendeley and are publicly available as of the date of publication. Accession links for both are listed in the key resources table. Microscopy data reported in this paper will be shared by the lead contact upon request.
- This paper does not report original code.
- Any additional information required to reanalyze the data reported in this work paper is available from the lead contact upon request.

### Experimental model and study participant details

- Mouse strains used as experimental models in this study:

Both male and female mice were used for breeding, whereas the phenotype (modeling breast cancer) was only explored in female mice. There is no phenotype present in male mice, hence no associations with sex could be recorded.

MMTV-rtTA (MTB)

*PyMT-IRES-Cre* (*MIC*)

*MMTV-NIC* (*NIC*)

*INSR^F/F^* (B6.129S4(FVB)-*Insr^tm1Khn^*/J; stock #006955)

*FVB/*NJ mice (stock#001800)

INSR^F/F^ FVB/NJ

- All mice used in this study were housed in the UHN animal facilities accredited by the Canadian Council on Animal Care and treated in accordance with the institutional guidelines and the UHN Animal Care Committee-approved protocol UHN AUP:2176.
- Cell lines used in this study:

MCF10A, human, female, from a mammary gland of a White, 36-year-old female with fibrocystic breasts.

Comma1D, mouse, female, from a mammary gland of 1-year-old female BALB/c mouse.

## Method details

### Gene expression analyses from published datasets of human samples and cell lines

RNA-seq gene expression data from the Cancer Cell Line Encyclopedia (Broad, 2019; “CCLE”) ^75^ dataset were obtained via the cBioPortal ^76,77^. Boxplots, scatterplots, and p-values of these data were visualized and calculated using GraphPad Prism 9.

### Culture of human cell lines

MCF-10A cells were cultured in DMEM/F12 (Invitrogen #11330-032) supplemented with 5% fetal bovine serum (FBS), 20 ng/mL epidermal growth factor, 0.5 mg/mL hydrocortisone, 100 ng/mL cholera toxin, 10 µg/mL insulin and 1% penicillin/streptomycin ^78^. HEK-293T, MCF-7 and MDA-MB-231 cells were cultured in DMEM (Invitrogen #D5796-500ML) supplemented with

10% FBS and 1% penicillin/streptomycin. For immunoblot signalling analyses, MCF-10A cells were seeded in above culture medium (Basal), followed by either basal medium replenishment or overnight starvation with DMEM/F12 supplemented with only 0.5 mg/mL hydrocortisone, 100 ng/mL cholera toxin and 1% penicillin/streptomycin (Starved). These cells were then stimulated with 10 µg/mL insulin in either the absence (Starved + Ins) or presence (Starved + OSI + Ins) of pre-treatment with 1 µM linsitinib (OSI-906) for 2 hours. For shRNA analyses, MCF-10A cells were transduced with lentiviral pLKO.1 puromycin shRNA vectors (RNAi Consortium, Broad Institute) to target expression of either *INSR* (shRNA 80 = 5’-GTG-CTG-TAT-GAA-GTG-AGT- TAT-3’, shRNA 82 = 5’-CTC-GTT-TGG-TTA-CCA-ATT-TAA-3’) or a non-targeting control (*shGFP*, 5’-GCA-AGC-TGA-CCC-TGA-AGT-TCA-T-3’), followed by growth in either Basal, Starved or Starved + Ins conditions outlined above.

### PCR of human breast cancer cell lines

Cells were grown under normal culture conditions followed by lysis with TRIzol Reagent (Invitrogen # 15596026), phase separation with chloroform, and RNA precipitation with isopropanol. RNA was then washed with 70% ethanol and re-dissolved with DEPC-treated water. First-strand cDNA synthesis from isolated mRNA was performed as outlined in the protocol for Superscript III Reverse Transcriptase (Invitrogen #18080051). For cDNA isolated from HEK-293T, MCF-10A, MCF-7 and MDA-MB-231 cells, polymerase chain reaction (PCR) was performed using primers spanning *INSR* exon 11 (Fwd: 5’-AAC-CAG-AGT-GAG-TAT-GAG-GAT-3’; Rev: 5’-CCG-TTC-CAG-AGC-GAA-GTG-CTT-3’) to specifically detect *INSR* isoforms as previously described ^79^. PCR products (*INSR-A* (exon 11-) = 600 bp; *INSR-B* (exon 11+) = 636 bp) were separated by agarose gel electrophoresis, allowing for relative *INSR* isoform expression levels to be calculated by densitometry analysis using ImageJ software, with visualization of relative *INSR* isoform expression performed in GraphPad Prism 9.

For shRNA knockdown experiments in MCF-10A cells, cells were lysed and first-strand cDNA synthesis was performed as described above, with real-time quantitative PCR (RT-qPCR) performed using primers targeting *INSR* (Fwd: 5’-AAC-CAG-AGT-GAG-TAT-GAG-GAT-3’; Rev: 5’-CGT-TCC-AGA-GCG-AAG-TGC-TT-3’) and *RPLPO* (Fwd: 5’- CAA-CCC-TGA-AGT-GCT-TGA-CAT-3’, Rev: 5’-AGG-CAG-ATG-GAT-CAG-CCA-3’). Fold change in mRNA expression between shRNA-treated and untreated cells was calculated using the 2^−ΔΔCq^ method as previously described ^80^ with *RPLPO* expression used as a reference for normalization. Calculated results were visualized in GraphPad Prism 9.

### Immunoblotting

To prepare total protein lysates, cells were homogenized in CHAPS lysis buffer (0.3% CHAPS, 120 mM NaCl, 1x protease inhibitor cocktail in 1.5 mM HEPES buffer pH 7.5). To detect relative protein and phosphorylated protein levels, immunoblots were probed with antibodies against INSR (Millipore Sigma #07-724), phospho-INSR/IGF1R (Y1135/1136, Cell Signaling #3024), Akt (Cell Signaling #4691), phospho-AKT (S^473^) (Cell Signaling #4058), ERK (Cell Signaling #9102), phospho-p44/42 MAPK (Erk1/2) (T^202^/Y^204^) (Cell Signaling #9101), S6 (Cell Signaling #2217), and phospho-S6 (S235/S236, Cell Signaling #). GAPDH (Santa Cruz #sc-25778) was probed for as a loading control.

### Conditional INSR deletion in *MTB:MIC* or *NIC* mouse models

The animals used in these experiments were housed in the UHN animal facilities accredited by the Canadian Council on Animal Care and treated in accordance with the institutional guidelines and the UHN Animal Care Committee-approved protocols. *MMTV-rtTA* (*MTB*) mice were a generous gift of Dr. Lewis A. Chodosh from the University of Pennsylvania School of Medicine ^40^. *PyMT-IRES-Cre* (*MIC*) mice were a generous gift of Dr. William Muller from the Rosalind and Morris Goodman Cancer Research Centre at McGill University ^41^. *MMTV-NIC* (*NIC*) mice were a generous gift of Dr. Giuseppina Ursini-Siegel from the Lady Davis Institute for Medical Research at McGill University ^42^. *INSR^F/F^* (B6.129S4(FVB)-*Insr^tm1Khn^*/J; stock #006955) and *FVB/NJ* mice (stock #001800) were purchased from the Jackson Laboratory. *INSR^F/F^*were backcrossed onto the *FVB/NJ* background for 6 generations. Mice were genotyped using the following primers: *MMTV-rtTA* (Fwd: 5’ ACC GTA CTC GTC AAT TCC AAG GG 3’, Rev: 5’ CGC CAT TAT TAC GAC AAG 3’), *PyMT-IRES-Cre* (Fwd: 5’ GGA AGC AAG TAC TTC ACA AGG G 3’, Rev: 5’ GGA AAG TCA CTA GGA GCA GGG 3’), *neu* (Fwd: 5′- CCT CTG ACG TCC ATC GTC TC-3′, Rev: 5′-CGG ATC TTC TGC TGC CGT CG-3′), and the conditional *INSR* allele (Fwd: 5’ GAT GTG CAC CCC ATG TCT G 3’, Rev: 5’ CTG AAT AGC TGA GAC CAC AG 3’). Regular chow-fed (#7012, Teklad Diets) *MMTV-rtTA/PyMT-IRES-Cre/INSR^+/+, F/+ or^ ^F/F^*(*MTB:MIC INSR* (+/+), (F/+) or (F/F)) mice were administered 2 mg/mL doxycycline hyclate (Sigma-Aldrich #D9891) in their drinking water at 10-12 weeks of age. *NIC INSR* (+/+), (F/+) or (F/F) mice on regular chow were monitored weekly for palpable tumours; upon detection, weekly caliper measurements were performed until a humane endpoint was reached (total tumour volume ≥ 1000 mm^3^), followed immediately by sacrifice. For both mouse models, mammary tumours, lungs, and other tissues were excised, weighed (tumours only), counted (tumours only) and fixed in 5% formalin for 48 hr at 4° C. Post-fixation, tumours and tissues were rinsed with PBS and stored in 70% ethanol at 4°C. Visualization and statistical analysis of all measurements were performed in GraphPad Prism 9.

### Blood glucose concentrations in INSR knockout mice

To ensure that INSR signaling was intact in the metabolic tissues of INSR knockout mice, their blood glucose and plasma insulin concentrations were measured. Mice were deprived of food for 6 hours, at which time blood glucose concentrations were measured using Accu-Chek Performa test strips and meter (Roche). Visualization and statistical analysis of all measurements were performed in GraphPad Prism 9.

### Statistical modeling and analysis of tumour growth data

Tumour growth data from *NIC INSR* mice was filtered to ensure that all tumours considered had at least two measurements before the mouse was sacrificed. Visual examination of the data suggested an exponential growth pattern for the tumours. To directly test for this, both linear and log-linear mixed models were fit to the measured tumour volume data over time to estimate the average tumour growth rate for each mouse. The measured volume of each tumour was modelled as a function of the tumour age, including fixed effect terms for the interaction between the genotype and the age of the tumour, as well as a fixed effect intercept for each genotype. Random effects were included for the mouse-to-mouse variation in the slope of the model, and for the tumour-to-tumour variation in the intercept. The model estimated an average tumour growth rate (linear or exponential) for each genotype, allowing for random variation in the growth rates between mice. The relationships between the residual and fitted values for the linear and log-linear models were visually examined, with the linear model displaying a pronounced dependency between the residuals and the fitted volume, suggesting a biased fit to the observed data, validating the use of the log-linear model. Estimated percent growth per day values were calculated from the coefficients estimated by the model for each mouse, adding together the average genotype growth rate and the random mouse-specific deviation from the average.

The significance of differences between groups was assessed using a nested-model ANOVA, comparing the full model described to a restricted model which excludes the fixed and random effects of genotype on growth rate (but retains the effect of genotype on the intercept). The tested null hypothesis was that a single average growth rate explains the observed data, as well as an average growth rate fit to each genotype separately. Two group comparisons were done by first refitting both models to the subset of the data belonging to the two compared groups.

The same data were also analysed after summing the tumour volumes for each mouse, calculating total tumour volume per mouse. Note that the previous analysis estimated an intercept term for each tumour, meaning the slopes were fit after adjusting for the different time of tumour onset. In per mouse analysis, the volume of tumours arising from the same mouse is added to the mouse total as they arise. For modelling these data, the models described above were modified to fit one random intercept per mouse. The same approach was taken for this analysis: after visual examination, linear and log-linear mixed models were fit to the data and compared based on a bias of residuals based on fitted total tumour volumes. As in per tumour analysis, the log-linear model was found to be a better fit than the linear model. Significance testing was done as above, comparing the explanatory power of models fit using the genotype information to those fitting a single slope for mice from all genotypes.

All models were estimated using the afex (1.0-1) (CRAN.R-project.org/package=afex) and lme4 (1.1-27.1) ^81^ packages, as implemented in the R Statistical Computing Environment (4.1.2) (<u>R-project.org/</u>). Plots were generated using the ggprism (1.0.3) (<u>CRAN.R-project.org/package=ggprism</u>) and ggplot2 (3.3.5) (<u>ggplot2.tidyverse.org</u>) packages.

### Immunohistochemistry and semi-automated digital image analysis

For histopathological and signaling analyses, FFPE tumours of *MTB:MIC INSR* and *NIC INSR* mice collected at endpoint were sectioned and stained with hematoxylin and eosin (H&E), or probed with antibodies against INSR (Upstate #05-1104), IGF1R (Cell Signalling #3027), HER2 (Santa Cruz #284), phospho-AKT (S^473^) (p-AKT; Cell Signaling #4058), phospho-p44/42 MAPK (Erk1/2) (T^202^/Y^204^) (p-ERK; Cell Signaling #9101), and Ki67 (Lab Vision #RM-9106-S1, clone SP6) by the Applied Molecular Profiling Laboratory and the Pathology Research Program at UHN. All slides were digitized for subsequent quantification by image analysis.

INSR, IGF1R, p-AKT, p-ERK and Ki67 stains were quantified using QuPath bioimage analysis software ^43^. Briefly, images of all slides of a given stain were loaded into a single project file where stain vector and background estimates for each antibody were used to improve stain separation using colour deconvolution. *Simple Tissue Detection* command was used to batch process the selection of tissue from tumour sections and exclusion of white space on all project slide images, followed by manual annotation of each tissue sample and exclusion of any stain artifact. The *Positive Cell Detection* command, which detects paired nuclei and cell boundaries within an annotation and scores each cell for staining intensity, was run with its parameters manually set and intensity thresholds optimized for each stain based upon mean nuclear (Ki67), cytoplasm (INSR, IGF1R, p-AKT) and cell (p-ERK) DAB optical densities of positive and negative control annotations. All project annotations were then batch processed with the optimized *Positive Cell Detection* parameters, followed by tissue classification using a Random Forest Classifier trained on manual annotations of tumour or stomal cells which finally enumerated tumour tissue-specific cells that were either negative (0 < lowest intensity threshold parameter) or positive (weak (1+) ≥ lowest intensity threshold parameter; moderate (2+) ≥ 2-fold 1+; strong (3+) ≥ 3-fold 1+) for a stain. Tumour % Positivity and Tumour H-scores for each annotation were quantified by QuPath, with subsequent visualization and statistical analysis performed in GraphPad Prism 9. Representative images of stained sections were visualized in QuPath at 20X magnification.

To validate tissue-specificity of the INSR knockout, FFPE liver, muscle and adipose (fat) tissues in *MTB:MIC INSR* (+/+) and (F/F) mice were sectioned, stained with anti-INSR and digitized for image visualization. To count metastases for both *MTB:MIC INSR* and *NIC INSR* mice, FFPE lungs collected at endpoint were serially sectioned (5 µm sections, 100 µm apart) and stained with H&E, followed by digitization and analysis using Aperio ImageScope (Leica Biosystems). The number of metastases were averaged across 5 sections, with subsequent visualization and statistical analysis performed in GraphPad Prism 9.

### Mammary tumour sphere forming assay

*In vitro* Matrigel colony forming assay was performed with adaptations from a method previously described ^82^. Mammary glands of 6-12 week old *FVB* (control) mice, *MTB:MIC INSR* (+/+),and *MTB:MIC INSR* (F/F) mice were resected, minced and digested in 4 ml of DMEM/F12 medium supplemented with 750 U/ml collagenase and 250 U/ml hyaluronidase for 1.5 hours at 37^0^C. Cell preparations were vortexed and mammary gland digestion halted with 10 mL of HF buffer (Hank’s Buffered Salt Solution supplemented with 2% FBS), followed by centrifugation at 1200 rpm for 5 minutes at 4^0^C. Red blood cells were lysed by resuspension of cell pellet in a 1:5 ratio of HF buffer to ammonium chloride, followed by centrifugation. Resulting cell pellet was gently resuspended for 2 minutes in 2 mL 0.25% trypsin, followed by dilution with HF buffer and centrifugation. Cell pellet was again gently resuspended for 2 minutes in 2 mL of 5 µg/mL dispase solution supplemented with 100 µL of 1 mg/mL DNAse I. After addition of 10 mL of HF buffer, cell suspension was filtered through a 40 µm cell strainer before centrifugation. Cell pellet was resuspended in HF buffer, and viable cell numbers determined using Trypan blue exclusion. 5000 cells were seeded per well in 50 µL Matrigel (Corning #356231) in 24 well plates and cultured in mouse Epicult-B medium (DMEM:F12 medium supplemented with 5 µg/mL insulin, 10 ng/mL EGF, 10 ng/ML cholera toxin, 180 µM adenine, 0.5 µg/mL hydrocortisone, 10% FBS, and 10 µM Rock inhibitor). Cells were treated daily with or without 0.1 µg/mL doxycycline hyclate for 11 days.

The Z-stack and montage functions of the Cytation 5 imaging multi-mode reader (BioTek) were used to image entire Matrigel domes containing mammary tumour spheres. The brightfield Z-stacks were converted to Z-projections and processed by digital phase contrast using the Gen5 software (Cytation 5). Tumour sphere morphometric data for each well was collected by performing image analysis on processed digital phase contrast images in Gen5. Briefly, all objects identified by Gen5 were filtered based on calculated metrics to identify all “spheres” in a well, and further filtering classified a subset of these as “tumour.” The parameters for the “tumour” filter were based on the following calculated metrics: size > 250; perimeter < 5000; circularity ≥ 0.2; mean intensity ≥ 25,000; eccentricity ≥ 0.1. Eccentricity (or “spikiness”) was used as a measure of sphere budding formation and was defined as: 1 – ((4π*Sphere Area)/(Sphere Perimeter^2^)). Visualization and statistical analysis of counts of mammary tumour spheres were performed in GraphPad Prism 9.

### INSR deletion by CRISPR/Cas9 in COMMA1D murine mammary epithelial cells

COMMA1D cells, a non-transformed murine mammary epithelial cell line, were cultured in COMMA1D media (DMEM/F12 media (Wisent Inc #319-005-CL) supplemented with 5% FBS (Wisent Inc #098150,), 20ng/ml EGF (Peprotech #AF-100-15), 0.5mg/ml Hydrocortisone (Sigma #H0888), 100ng/ml Cholera Toxin (Sigma C8052), 10ug/ml Insulin (Biogems #10-365), and 1% Penicillin/Streptomycin (Wisent Inc #450-201-EL)). Cells were stained with FITC anti-mouse Ly- 6A/E (Sca-1; Biolegend #122504) and APC/Cy7 anti-mouse CD326 (Ep-CAM; Biolegend #118217) and sorted by FACS (Aria Fusion, BD Biosciences) to isolate the luminal (Ep-CAM high:Sca-1 low) and basal (Ep-CAM low:Sca-1 high) populations ^46^. Sorted luminal COMMA1D cells were then genetically modified to delete *INSR* using CRISPR/Cas9 as described in ^44^. Briefly, px458 (Addgene #62988), a plasmid containing a Cas9n D10A nickase mutant with a puromycin selection marker, was cloned to incorporate knock-out sgRNAs targeting *INSR* (Top: 5’-CAC-CGG-GTA-TAA-GTC-TCT-CAT-TTG-G-3’, Bottom: 5’-AAA-CCC-AAA-TGA-GAG-ACT-TAT-ACC-C-3’). COMMA1D cells were then transfected with *INSR* knock-out plasmid using 15 µL PolyJet transfection reagent (SignaGen Laboratories #SL100688). Following transfection, cells were selected with 5 µg/ml puromycin (Sigma #P8833) for 48 hours, after which selection pressure was removed and cells allowed to proliferate until colonies formed (∼4 days). Clones were picked, expanded in COMMA1D media, and screened for deletion of *INSR* by PCR (Fwd primer: 5’-CTC-ATG-TGC-AGC-TAG-CTT-TC-3’, Rev primer: 5’-GAG-ATT-CCT-GCT-GCA-GAT-TG-3’). Two COMMA1D *INSR* knock-out (-/-) clones (#87, #160) and two control COMMA1D *INSR* wild-type (+/+) clones (#44, #65) were selected, expanded in COMMA1D media, and validated for INSR knock-out or expression by immunoblotting. Immunoblot analysis of canonical INSR signaling was performed as described above. Analysis of immunoblot signal by densitometry was performed in ImageJ, with visualization and statistical analysis performed in GraphPad Prism 9.

### RNAseq of COMMA1D INSR cells

Cells derived from two clones each of COMMA1D INSR (+/+) (Clones #44 and #65) and COMMA1D INSR (-/-) (Clones #87 and #160) were cultured in triplicate using COMMA1D media in 10 cm plates. RNA was then isolated using the RNeasy mini kit (Qiagen #74104) according to the manufacturer’s protocol and using the RNase free DNase set. 2 µg of purified RNA (each) from the triplicate samples was then sent to Novogene Inc. for RNAseq analysis.

Paired-end RNAseq reads were trimmed for Illumina adapters and sequencing quality using Trim Galore! (v. 0.6.6; https://www.bioinformatics.babraham.ac.uk/projects/trim_galore/) and aligned to the mm10 genome using STAR (v. 2.7.9a) ^83^. High-quality, single-mapping reads were counted to gene exons using FeatureCounts (subread v. 2.0.1) ^84^ and normalized and analyzed for differential expression between *INSR* wild-type (+/+) and knock-out (-/-) conditions using DESeq2 (v. 1.36.0) ^85^. Gene ontology enrichment analysis was performed using the Bioconductor package gprofiler2 (v. 0.2.1)^86^ and gene set enrichment analysis was performed using the Bioconductor package fgsea (v. 1.22.0) ^87^ and manually curated gene sets from http://download.baderlab.org/EM_Genesets/November_17_2022/Mouse/symbol/Mouse_GOBP_AllPathways_no_GO_iea_November_17_2022_symbol.gmt. Gene ontology cellular component gene sets related to mitochondrial respiratory chain complexes were obtained from g:profiler gene sets.

### Analysis of published RNAseq data from mouse preadipocytes

Raw RNAseq gene-level counts for INSR/IGF1R-depleted cells reconstituted with various forms of human INSR were obtained from GEO (Accession: GSE206565) ^47^. Counts were normalized using DESeq2 and mitochondrial respiratory chain complex gene sets were applied to z-score scaled gene expression.

### Analysis of INSR CPTAC data

Relative protein abundance values (TMT log2 ratio) for INSR from breast cancer samples and adjacent normal samples were downloaded from the Cancer Proteogenomic Data Analysis Site (cprosite.ccr.cancer.gov) and were paired with clinical metadata obtained from the CPTAC Prospective Breast BI Proteome project (pdc.cancer.gov/pdc; file: S039_BRCA_prospective_clinical_data_r2.xlsx)^36^. Relative protein abundance was compared between tumor and normal adjacent samples as well as estrogen receptor/progesterone receptor positive/negative samples, HER2 positive/negative samples, and triple negative breast cancer (TNBC) samples and samples not categorized as TNBC using Wilcoxon rank-sum tests.

### RT-qPCR of murine mammary epithelial cells

Total RNA from COMMA1D INSR (+/+) (Clones #44 and #65) and COMMA1D INSR (-/-) (Clones #87 and #160) cultured in triplicate in COMMA1D media was isolated using the Qiagen RNeasy mini kit (Qiagen # 74104) according to the manufacturer’s protocol. First-strand cDNA synthesis from isolated mRNA was performed as outlined in the protocol for Superscript III Reverse Transcriptase (Invitrogen #18080051). Real-time quantitative PCR (RT-qPCR) was performed using the BioRad SsoAdvanced Universal SYBR green Supermix according to the manufacterer’s protocol (BioRad #172571). Briefly, 12.5ng cDNA was added to 5ul SsoAdvanced Universal SYBR Green Supermix in a skirted 96 well plate (BioRad #HSP9601) with the following primers: *INSR* (Fwd: 5’-AGA-TGA-GAG-GTG-CAG-TGT-GGC-T-3’, Rev: 5- GGT-TCC-TTT-GGC-TCT-TGC-CAC-A-3’), *ATP5b* (Fwd: 5’-CTC-TGA-CTG-GTT-TGA-CCG-TTG-C-3’, Rev: 5’-TGG-TAG-CCT-ACA-GCA-GAA-GGG-A-3’), *SHDA* (Fwd: 5’-GAG-ATA-CGC-ACC-TGT-TGC-CAA-G-3’, Rev: 5’-GGT-AGA-CGT-GAT-CTT-TCT-CAG-GG-3’), *UQCRB* (Fwd: 5’-CCA-TAA-GAA-GGC-TTC-CTG-AGG-AC-3’, Rev: 5’-TTT-GTC-CAC-TGA-TCC-TTA-GGC-AAG-3’), *Cox7c* (Fwd: 5’-ATG-TTG-GGC-CAG-AGT-ATC-CG-3’, Rev: 5’-ACC-CAG-ATC-CAA-AGT-ACA-CGG-3’), *Ndusf6* (Fwd: 5’-CGG-GGA-AAA-GAT-CAC-GCA-TAC-C-3’, Rev: 5’- TCC-ACC-TCG-TTC-ACA-GGC-TGT-T-3’), and *HPRT* (Fwd: 5’-CTG-GTG-AAA-AGG-ACC- TCT-CGA-AG-3’, Rev: 5’-CCA-GTT-TCA-CTA-ATG-ACA-CAA-ACG-3’). The RT-qPCR amplification was performed in the CFX Connect PCR machine (BioRad) using the CFX Manager 3.0 software (BioRad) with the following parameters: 39 cycles of 98^0^C for 5 seconds followed by 60^0^C for 20 seconds. Data was exported and mRNA expression was calculated using the 2^−ΔΔCq^ method as previously described ^80^. Calculated expression levels of *INSR*, *ATP5b*, *SHDA*, *UQCRB*, *Cox7c*, and *Ndusf6* were each normalized to mRNA expression of housekeeping gene *HPRT*. Visualization and statistical analysis of relative gene expression was performed in GraphPad Prism 9.

### Determination of relative mtDNA content in murine mammary epithelial cells

Total cellular DNA of COMMA1D INSR (+/+) (Clones #44 and #65) and COMMA1D INSR (-/-) (Clones #87 and #160) was extracted using the Qiagen DNeasy Blood and Tissue kit (Qiagen #69504). Relative mitochondrial DNA content was evaluated by qPCR targeting the β- globin gene and the mitochondrial *COX2* (*MT-COX2*) as previously described ^88^ using the following primers: *MT-COX2* (Fwd: 5’-GCC-GAC-TAA-ATC-AAG-CAA-CA-3’, Rev: 5’-CAA- TGG-GCA-TAA-AGC-TAT-GG-3’) and β-globin (Fwd: 5’-GAA-GCG-ATT-CTA-GGG-AGC-AG-3’, Rev: 5’-GGA-GCA-GCG-ATT-CTG-AGT-AGA-3’). The reaction was conducted in a 10 uL system containing 12.5ng DNA, 5ul SsoAdvanced Universal SYBR Green Supermix (BioRad #172571) in a skirted 96 well plate (BioRad #HSP9601), and 2 nM forward and reverse primers. The PCR amplification was performed in the CFX Connect PCR machine (BioRad) using the CFX Manager 3.0 software (BioRad) with the following process: 39 cycles of 98 ^0^C for 5 seconds followed by 60 ^0^C for 20 seconds. The qPCR results were calculated using the 2^−ΔΔCq^ method as previously described ^80^, and *MT-COX2* content was normalized to that of genomic β- globin. Visualization and statistical analysis of relative mtDNA content was performed in GraphPad Prism 9.

### Seahorse Mito Stress Test assay

Cells derived from two clones each of COMMA1D INSR (+/+) (Clones #44 and #65) and COMMA1D INSR (-/-) (Clones #87 and #160) cultured in COMMA1D media were seeded in 22 wells of a Seahorse 96-well plate (Agilent #103792-100) at a density of 15,000 cells per well. The following day, cells were replenished with XF-Base media (Agilent #103334-100) containing 10 mM glucose, 1 mM pyruvate, and 2 mM glutamine. The following compounds were loaded into the Seahorse XFe 96 (Agilent #103792-100): Port A, 1 µM oligomycin (Sigma #O4876); Port B, 2 µM FCCP (Sigma #C2920); Port C, 1 µM rotenone/Antimycin A (Sigma #R8875 and #A8674); and Port D, 20 µM monensin (Sigma #M5273). Following calibration, culture plates containing wild-type and knock-out COMMA1D INSR cells were introduced into the Seahorse XFe 96 tray and the acquisition program was run. This program included an equilibration step followed by three measurements of basal oxygen consumption rates (OCR) and extracellular acidification rates (ECAR), as well as three OCR and ECAR measurements each post-oligomycin injection, post-FCCP injection, post-rotenone/Antimycin A injection, and post-monensin injection. Readings were collected approximately 5, 10, and 15 minutes following the initiation of each step. Data were collected and analysed using the Wave 2.6 software (Agilent Technologies), with calculations of ATP production rates by mitochondrial respiration (J-ATPox) and glycolysis(J-ATPglyc) under basal, maximal OXPHOS, and maximal glycolytic conditions performed as previously described ^89^. Data visualization and statistical analyses were performed in GraphPad Prism 9.

### Glucose/galactose growth assay

COMMA1D INSR (+/+) and COMMA1D INSR (-/-) cells were cultured in COMMA1D glucose-free media (glucose-free DMEM/F12 media (Wisent Inc #319-176-CL) supplemented with 5% FBS (Wisent Inc #098150), 20ng/ml EGF (Peprotech #AF-100-15), 0.5mg/ml Hydrocortisone (Sigma #H0888), 100ng/ml Cholera Toxin (Sigma #C8052), 10ug/ml Insulin (Biogems #10-365), 1% Penicillin/Streptomycin (Wisent Inc #450-201-EL) additionally supplemented with 17 mM glucose (Sigma #G5767). Cells were counted using Trypan blue exclusion, followed by seeding of 200,000 cells each in 4 wells of two 6-well plates (performed in triplicate). The following day (Day 0), cells from one well each of the duplicate plates were washed in 1X PBS (Wisent Inc #311-010-CL), treated with 0.5 mL 0.25% trypsin, and viable cell numbers were determined using Trypan blue exclusion. Media was then removed from the remaining wells and cells were washed in 1X PBS, followed by replenishment in COMMA1D glucose-free media supplemented with either 17 mM glucose or 17 mM galactose (Sigma #G0750). Viable cell numbers were then determined as described above from one of the remaining wells of each duplicate plate at 24, 48, and 72 hours following Day 0 replenishment with either glucose- or galactose-containing media. Visualization and statistical analysis of viable cell counts were performed in GraphPad Prism 9.

### Expression of INSR and INSR KD in Comma1D cells

The human Insulin Receptor cDNA was a gift from Frederick Stanley (Addgene plasmid # 24049; http://n2t.net/addgene:24049; RRID:Addgene_24049). This construct (hIR) was mutated to a kinase dead INSR by changing amino acid 1030 from lysine to arginine. This was done by purchasing a BstEII to PmlI synthetic DNA fragment from TWIST Biosciences that contains the mutation described and then subcloning it into the above plasmid to create hINSR kinase dead (hINSR KD). The cDNA’s were *in vitro* transcribed from the T7 promoter using the mMESSAGE mMACHINE T7 Ultra kit from ThermoFisher Scientific according to the manufacturers protocol (#AM1345). The transcribed mRNA was cleaned up with the MEGAclear Kit (ThermoFisher Scientific, AM1908). The mRNAs were transfected into Comma1D luminal INSR WT and KO cells using Lipofectamine MessengerMAX Reagent according to the manufacturers protocol (ThermoFisher Scientific, LMRNA001). The Seahorse assay was performed 24hrs post transfection.

## Quantification and statistical analysis

GraphPad Prism 9 and R Statistical Computing Environment (4.1.2) statistical software were used to calculate statistical values. Details of statistical tests and definitions of n can be found in each corresponding figure legend. Briefly, for mouse tumour quantification, death and immune cell tumour infiltration, boxplots show the median and interquartile range, with Tukey style whiskers and scatterplots show the median and interquartile range. ** p < 0.01, *** p < 0.001, **** p < 0.0001. Log-rank test was used for inverse Kaplan-Meier curves, and Student’s t-test for tumour numbers. For QPath analysis, scatterplots show the median and interquartile range. * p < 0.05, ** p < 0.01, *** p < 0.001, **** p < 0.0001, n.s. = not significant; Student’s t-test. For RNAseq analysis, FDR<0.05 was used. RNAseq boxplots show the median and interquartile range, with Tukey style whiskers. * p < 0.05; ** p < 0.01; **** p < 0.0001 with Benjamini-Hochberg corrected Wilcoxon rank-sum test. For mRNA analysis by Q-PCR, bar graphs show the mean with error bars representing SEM. For Seahorse analysis bar graphs show mean with error bars representing SEM. Time-resolved plots show mean with error bars representing SEM. * p < 0.05, ** p < 0.01, *** p < 0.001, **** p < 0.0001, n.s. = non-significant; Student’s t-test. For INSR protein data analysis from CPTAC, p-values shown are from Wilcoxon rank-sum test. For western blot quantifications time-resolved plots of mean values are presented with error bars representing SEM. n.s. = non-significant; Student’s t-test.

### Additional resources

Not applicable.

## Supplemental Information

**Supplementary Figure 1:**
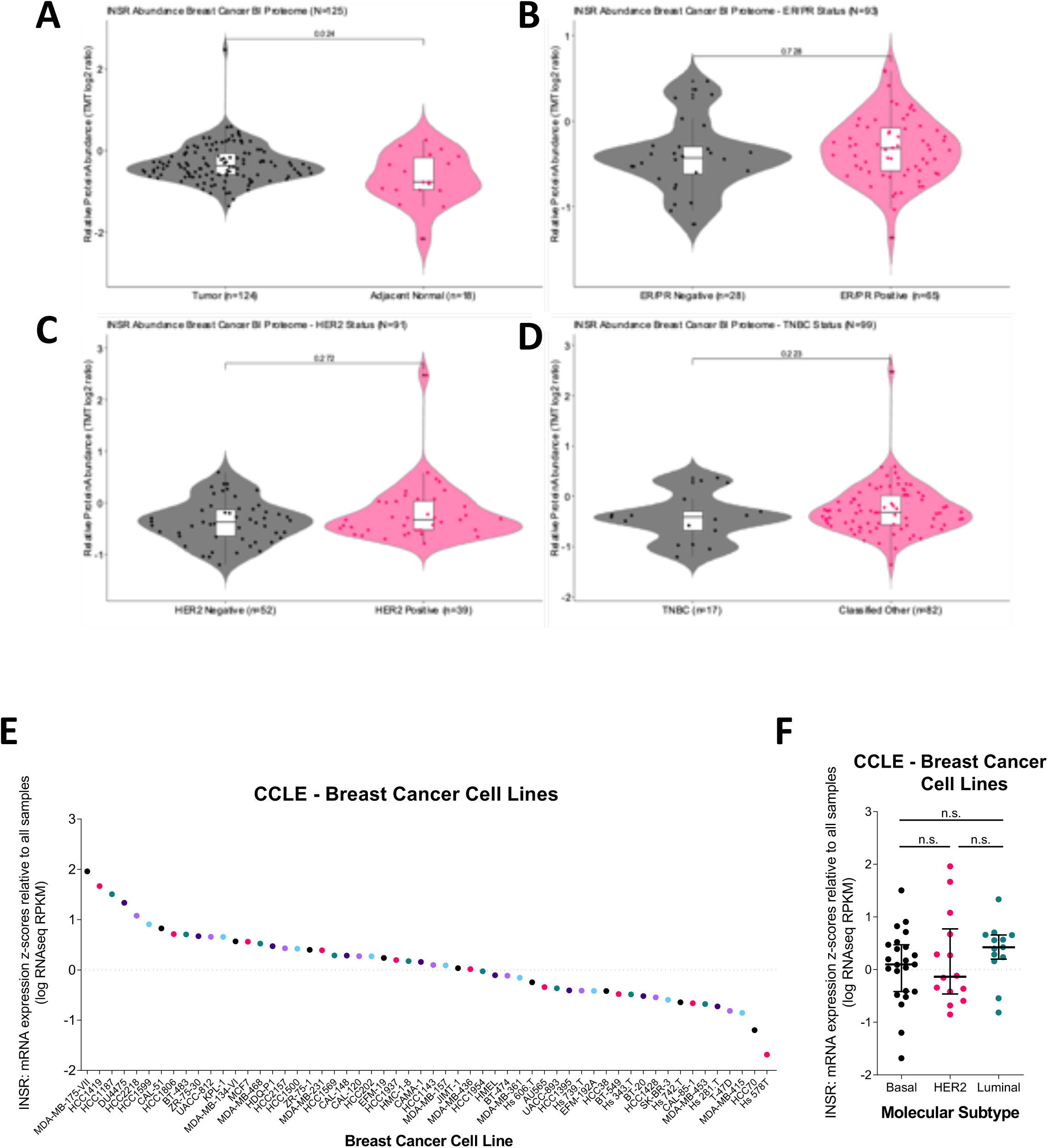
INSR protein abundance in breast cancer samples and INSR mRNA expression across breast cancer cell lines. (**A**) Violin plot showing the relative protein abundance of INSR between breast cancer samples (black; n=124) and adjacent normal tumors (pink; n=18). (**B**) Violin plot showing the relative protein abundance of INSR between breast cancer samples that are negative for estrogen receptor (ER) and progesterone receptor (PR) (black; n= 28) and samples that are ER/PR positive (pink; n= 65). (**C**) Violin plot showing the relative protein abundance of INSR between breast cancer samples that are negative for HER2 (black; n= 52) and samples that are positive for HER2 (pink; n= 39). (**D**) Violin plot showing the relative protein abundance of INSR between breast cancer samples that are classified as triple negative (TNBC; black; n= 17), and other breast cancer samples that were classified otherwise (Classified Other; pink; n= 82). p-values shown are from Wilcoxon rank-sum test. (**E**) Scatterplot of RNAseq expression analysis of *INSR* in breast cancer cell lines from the CCLE dataset found in cBioPortal. (**F**) Scatterplot of *INSR* mRNA expression in BC cell lines, grouped by molecular subtype, from the Cancer Cell Line Encyclopedia (“CCLE”) ^75^ dataset in cBioPortal.

**Supplementary Figure 2:**
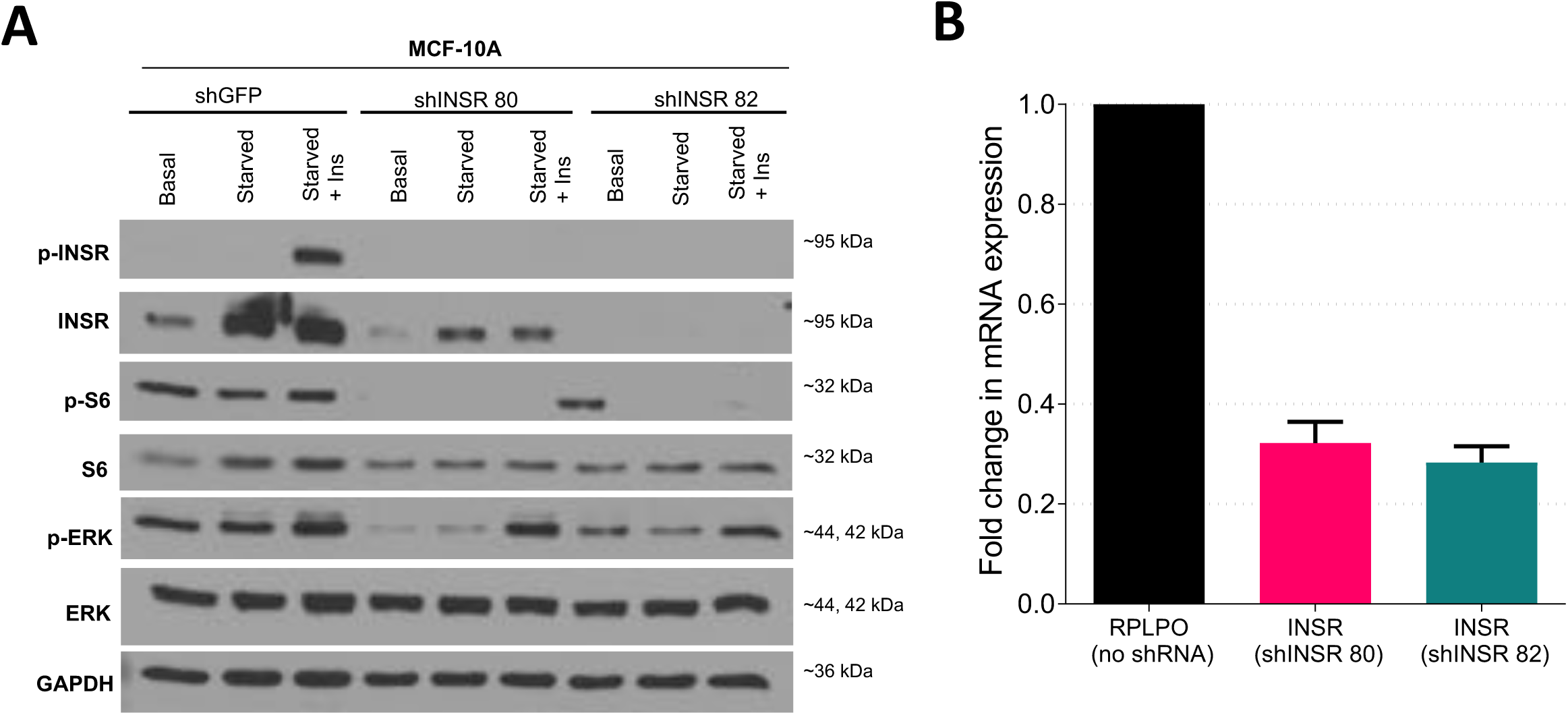
INSR signalling in mammary epithelial cells. (**A**) Western blot of MCF-10A cells transduced with either two shRNAs against *INSR* or a non-targeting control (*shGFP*), followed by growth in either supplemented DMEM/HAM (Basal) or starved overnight in the absence of insulin (Starved), and then stimulated with 100 nM insulin for 5 min (Starved + Ins). Cell signaling was analyzed using antibodies against p-INSR/p-IGF1R, INSR, p-ERK, ERK, p-S6, S6 and GAPDH and performed on 25 µg total cell protein. (**B**) Bar graph showing quantification of change in mRNA expression of *INSR* after treatment of MCF-10A cells with indicated shRNA against *INSR*, relative to *INSR* mRNA expression in MCF-10A cells treated with shGFP, analyzed via qRT-PCR. Change in *RPLPO* expression was used as a reference. Bar graphs show the mean with error bars representing SEM.

**Supplementary Figure 3:**
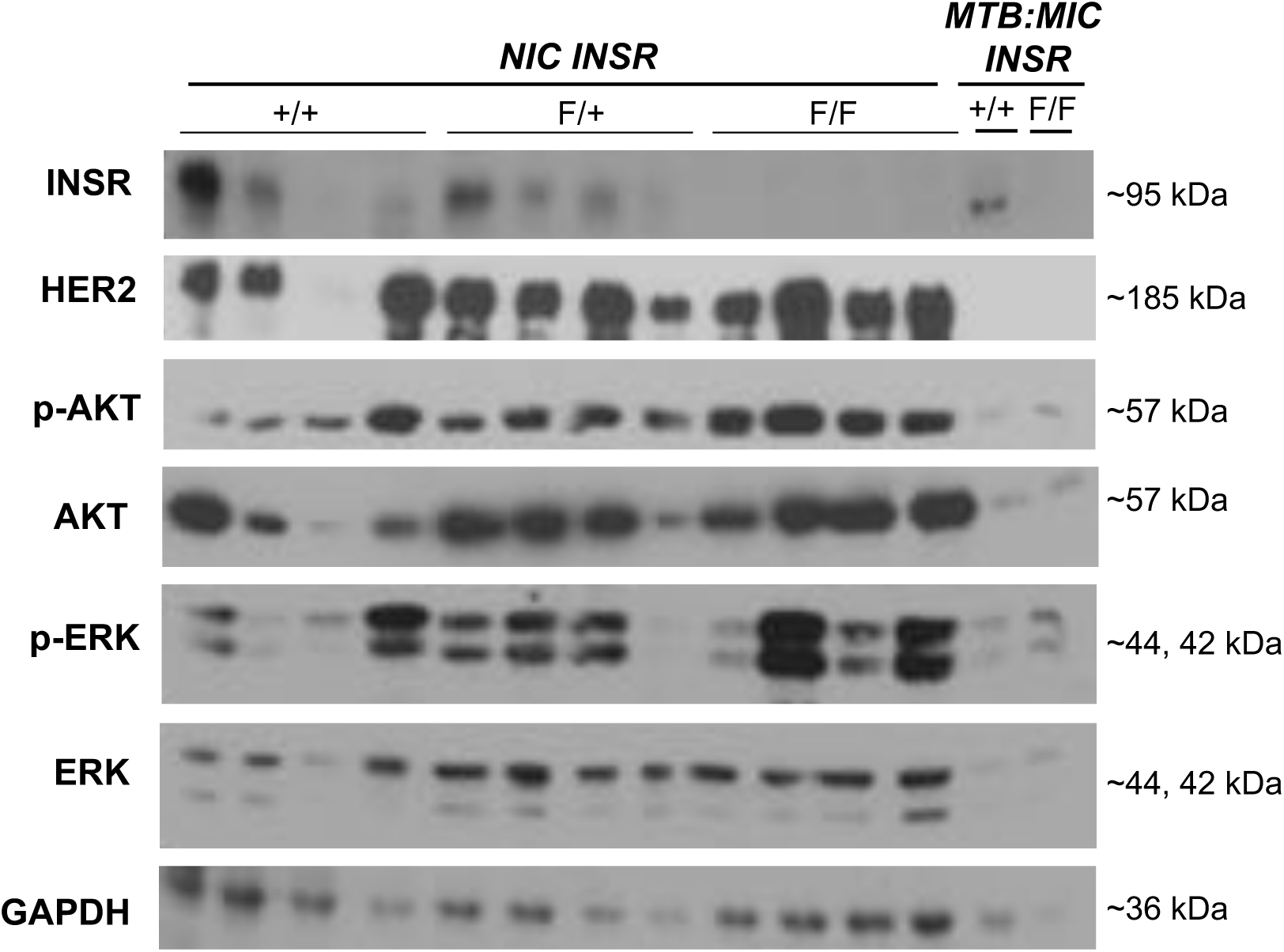
Immunoblotting analysis of NIC INSR and MTB:MIC INSR tumour lysates. Western blot analysis of lysates of *NIC INSR* (+/+, n = 4; F/+, n = 4; F/F, n = 4) and *MTB:MIC INSR* (+/+, n = 1; F/F, n = 1) mammary tumours snap frozen at endpoint. Blots were probed with antibodies to determine total levels of INSR, HER2, AKT and ERK, and phosphorylated levels of AKT and ERK. GAPDH was probed for as a loading control.

**Supplementary Figure 4:**
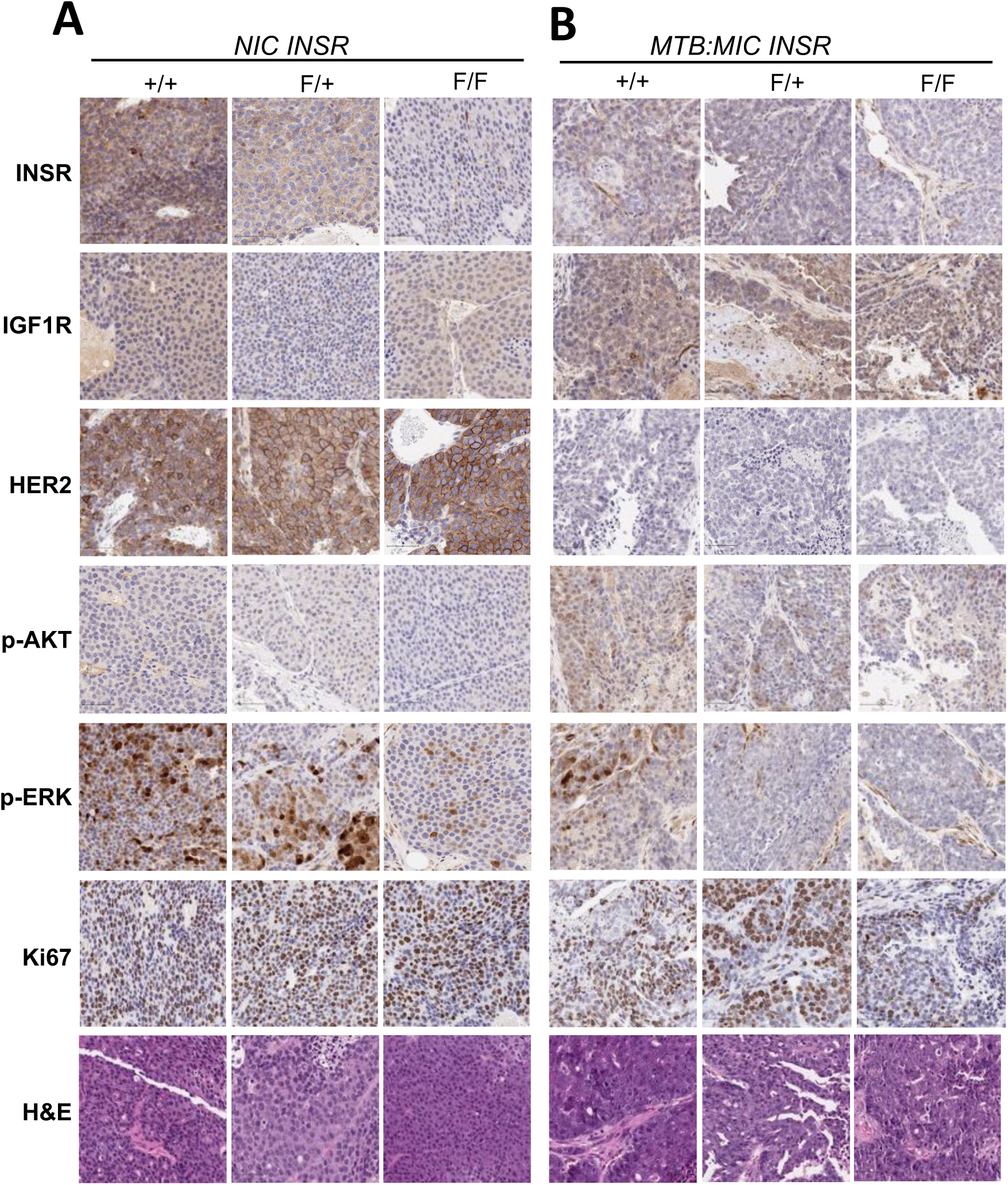
Immunohistochemistry of NIC INSR and MTB:MIC INSR tumour sections. Representative images from digital scans of INSR, IGF1R, HER2, p-AKT, p-ERK, Ki67 or H&E stained sections from endpoint tumours of (**A**) *NIC INSR* mice, or (**B**) *MTB:MIC INSR* mice. Images represent 20X magnification.

**Supplementary Figure 5:**
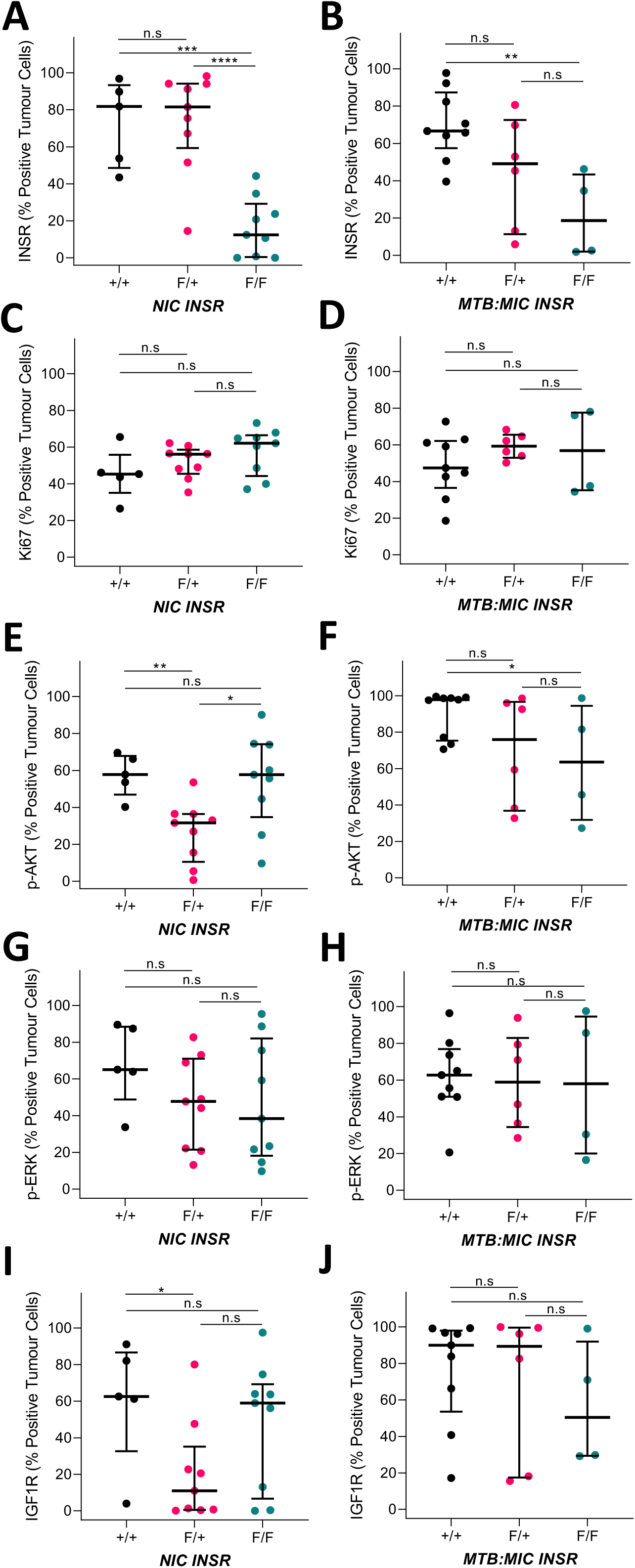
Semi-automated digital image analysis of NIC and MTB:MIC tumour sections. Scatterplots of digital pathological analysis using QuPath software of immunohistochemically stained sections of paraffin embedded tumours collected at endpoint from (A, C, E, G, I) *NIC INSR* mice (+/+, n = 5; F/+, n = 9; F/F, n = 9), (B, D, F, H, J) *MTB:MIC INSR* mice (+/+, n = 9; F/+, n = 6; F/F, n = 4). The percentage of positive tumour cells (% Positive Tumour Cells) in scanned images of stained tumour sections were determined by QuPath using *Simple Tissue Detection* followed by *Positive Cell Detection* and tissue classification, which selectively scored individual tumour cells as positive (1+, 2+ or 3+) or negative (0) for staining of (A-B) INSR, (C-D) Ki67, (E-F) p- AKT, (G-H) p-ERK, and (I-J) IGF1R. Scatterplots show the median and interquartile range. * p < 0.05, ** p < 0.01, *** p < 0.001, **** p < 0.0001, n.s. = not significant; Student’s t-test.

**Supplementary Figure 6:**
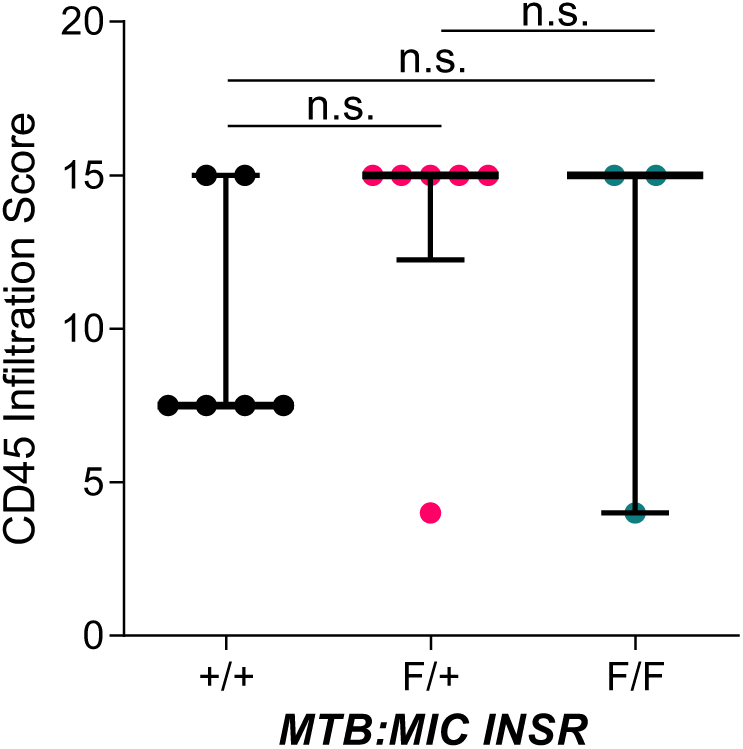
INSR loss does not impact immune infiltration in progressing mammary tumours. Scatterplot of CD45 infiltration score of tumours collected from *MTB:MIC INSR* mice (+/+, n = 6; F/+, n = 6; F/F, n = 3) following 10-week doxycycline induction. Scatterplot shows the median and interquartile range. n.s. = not significant; Student’s t-test.

**Supplementary Figure 7:**
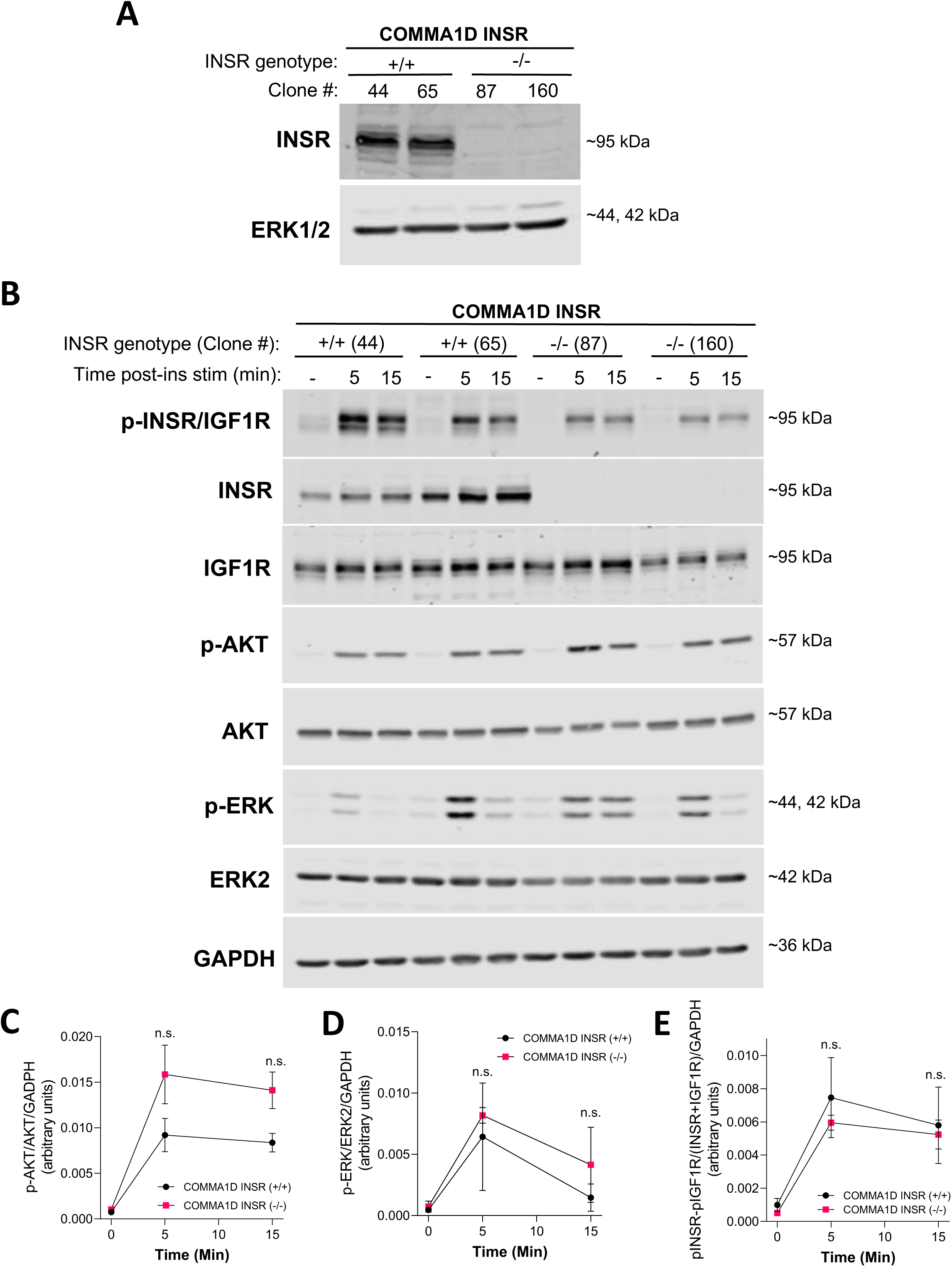
INSR loss in murine mammary epithelial cells does not impact canonical INSR signaling. Murine mammary epithelial cell line COMMA1D cells were sorted by FACS to isolate luminal population, which were then transfected with plasmids containing *Cas9* and sgRNAs designed to knock out *INSR*. (**A**) Western blot analysis of lysates of COMMA1D INSR wild-type (+/+) (n = 2) and COMMA1D INSR knockout (-/-) (n = 2) clones. Blots were probed with antibody against INSR. ERK1/2 was probed for as a loading control. (**B**) Western blot analysis of lysates of parental COMMA1D cells (n = 1) as well as COMMA1D INSR (+/+) (n = 2) and COMMA1D INSR (-/-) (n = 2) clones starved overnight, followed by stimulation with 100 nM insulin (ins). Lysates were collected immediately prior to stimulation (-), as well as 5 and 15 minutes post stimulation. Blots were probed with antibodies against p-INSR/IGF1R, INSR, IGF1R, p- AKT, AKT, p-ERK, and ERK2. GAPDH was probed as a loading control. (**C-E**) Time-resolved plots quantifying the relative optical density of the (**C**) p-AKT (normalized to AKT), (**D**) p-ERK (normalized to ERK2), and (**E**) p-INSR/IGF1R signal (normalized to the sum of INSR and IGF1R) of COMMA1D INSR (+/+) (n = 2) clones and COMMA1D INSR (-/-) (n = 2) clones, as measured by ImageJ software. Values were normalized to the optical density of GAPDH. Time-resolved plots show the mean value with error bars representing SEM. n.s. = non-significant; Student’s t-test.

**Supplementary Figure 8:**
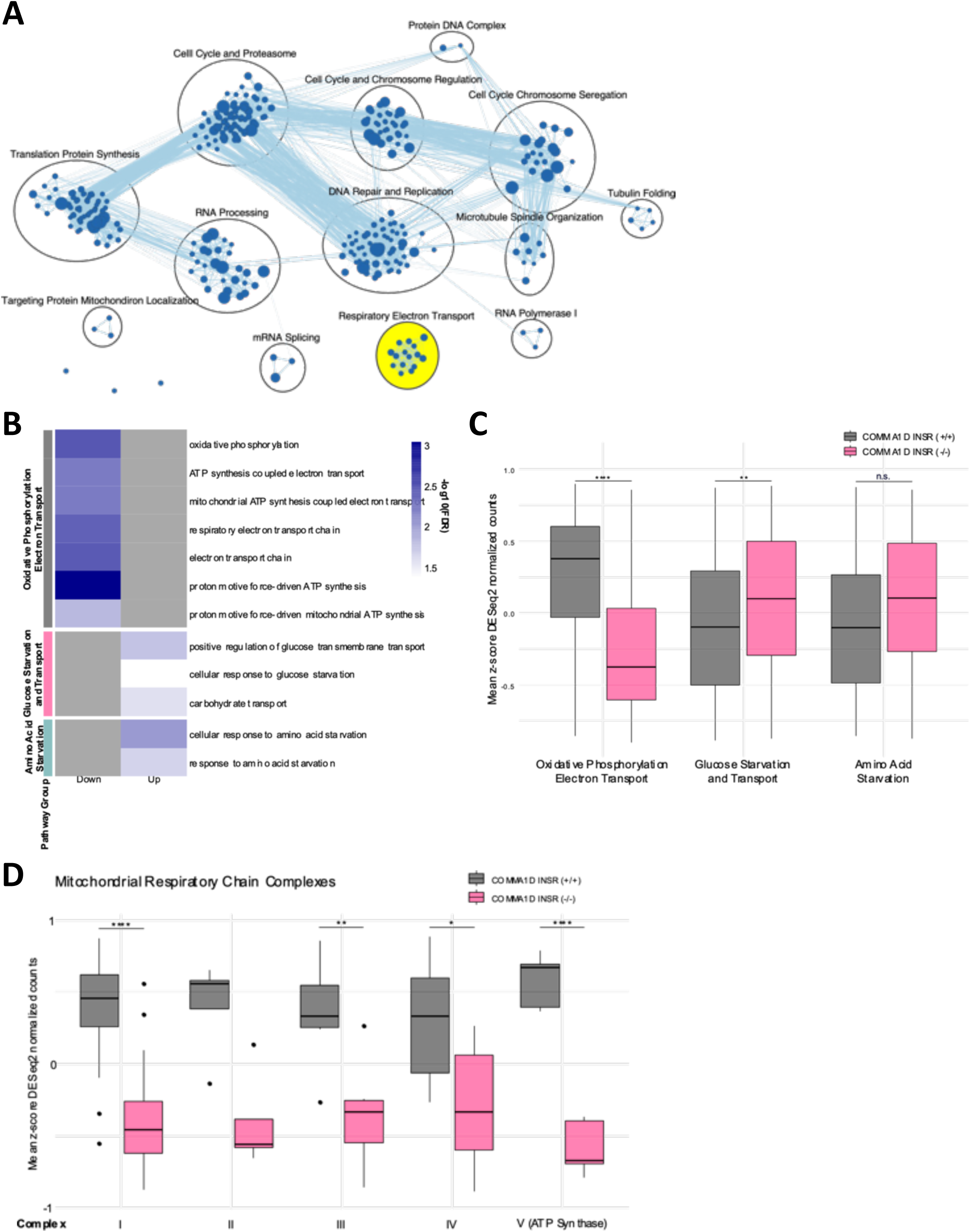
Loss of INSR in mammary epithelial cells reduces expression of mitochondrial respiratory chain complex components. (**A**) Enrichment map generated in Cytoscape of 358 significantly enriched pathways from gene set enrichment analysis (GSEA; FDR < 0.1) of COMMA1D INSR (-/-) cells (n = 6), relative to COMMA1D INSR (+/+) cells (n = 6), as measured by RNAseq. Each node represents a pathway and each edge represents an overlap of shared genes between pathways of 0.5. All pathways in this analysis are downregulated in COMMA1D INSR (-/-). Related pathways are clustered. (**B**) Heatmap showing significance [-log10(FDR)] of gene ontology biological processes associated with oxidative phosphorylation/electron transport, glucose starvation and transport, and amino acid starvation enriched in down- or up-regulated genes differentially expressed between COMMA1D INSR (-/-) (n = 6) and COMMA1D INSR (+/+) (n = 6) cells, as measured by RNAseq. Dark-grey cells indicate no significant enrichment. (**C**) Boxplot of mean z-score normalized counts by RNAseq for COMMA1D INSR (+/+) (n = 6) and COMMA1D INSR (-/-) (n = 6) cells for genes associated with biological processes related to oxidative phosphorylation/electron transport (# of genes: 183), glucose starvation and transport (# of genes: 199), and amino acid starvation (# of genes: 51). (**D**) Boxplot of mean z-score normalized counts by RNAseq for COMMA1D INSR (+/+) cells (n = 6) and COMMA1D INSR (-/-) cells (n = 6) for genes associated with mitochondrial respiratory chain complexes. Boxplots show the median and interquartile range, with Tukey style whiskers. * p < 0.05; ** p < 0.01; **** p < 0.0001; Benjamini-Hochberg corrected Wilcoxon rank-sum test.

**Supplementary Figure 9:**
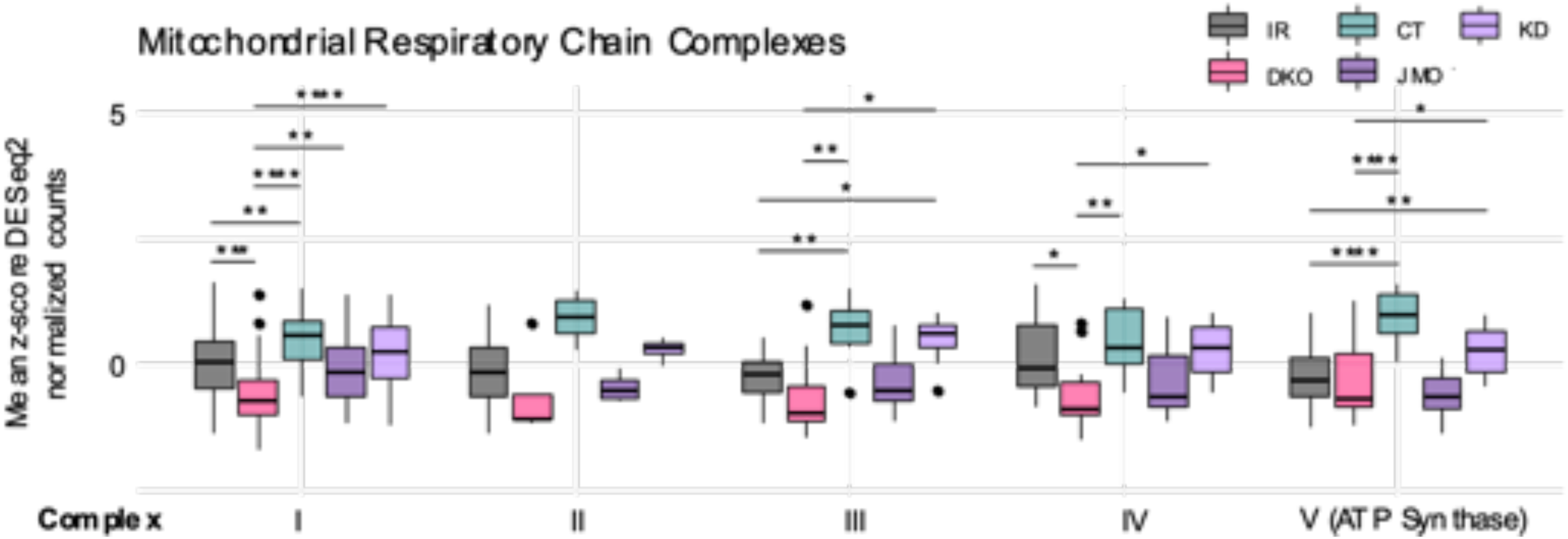
Impact of INSR and IGF1R loss on mitochondrial respiratory chain subunit expression in mouse preadipocytes is rescued by kinase-dead INSR. Boxplot of mean z-score normalized expression by RNAseq from GSE206565 ^47^ in mouse preadipocytes with a double knockout for INSR and IGF1R (DKO; n = 3) and DKO cells expressing a rescue with the wild-type human INSR gene (IR; n = 3), a truncated human INSR missing the C-terminus (CT; n = 3), a human INSR juxtamembrane domain only (JMO; n = 3), and a kinase-dead human INSR (KD; n = 3). Genes shown are associated with mitochondrial respiratory chain complexes. Boxplots show the median and interquartile range, with Tukey style whiskers. * p < 0.05; **p < 0.01; ***p < 0.001; **** p < 0.0001; Benjamini-Hochberg corrected Wilcoxon rank-sum test.

**Supplementary Figure 10:**
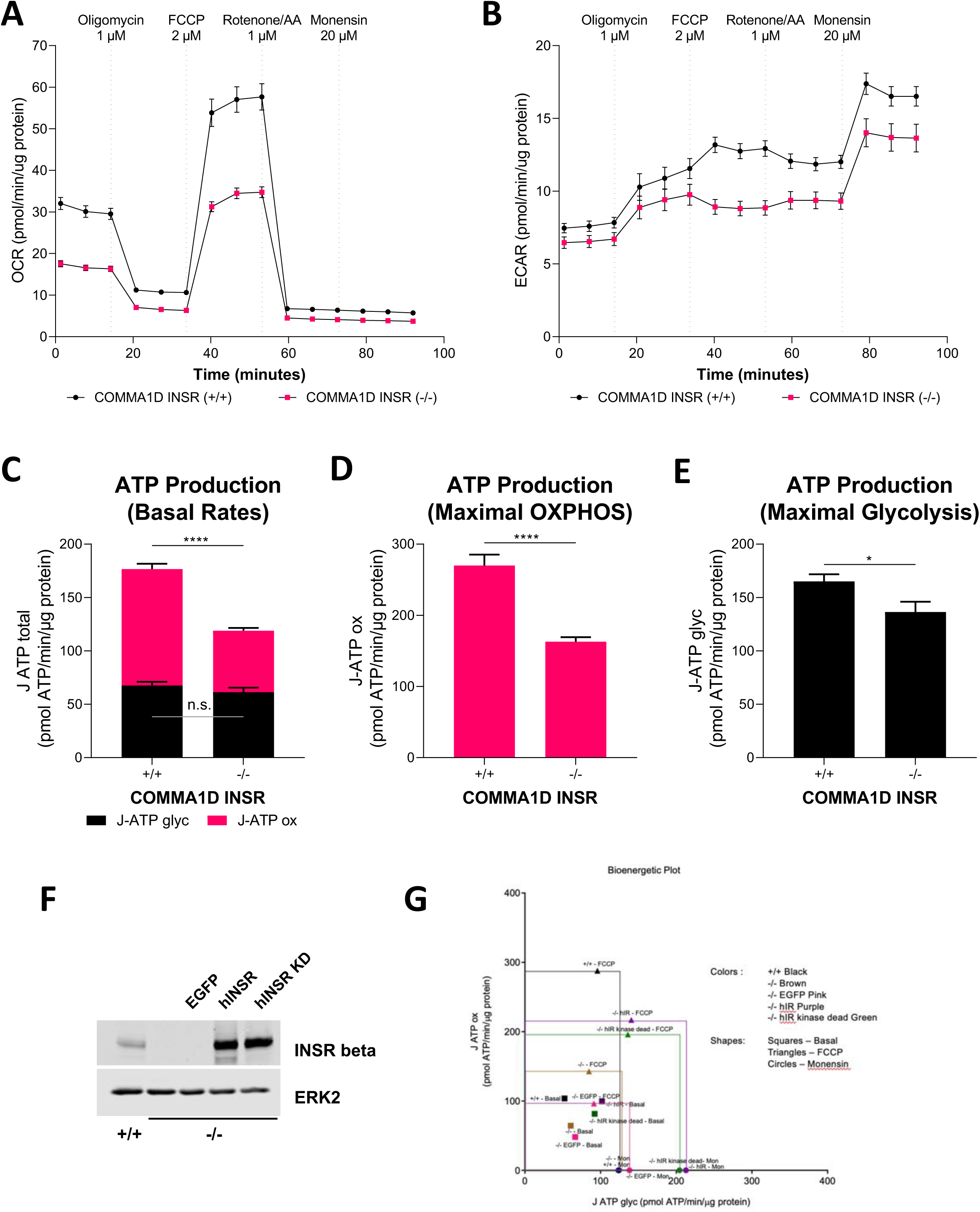
Impact of INSR loss INSR reexpression on bioenergetic fitness in mammary epithelial cells. Seahorse Mito Stress Test Assay of COMMA1D INSR (+/+) (n = 39) and COMMA1D INSR (-/-) (n = 40) cells seeded onto 96-well Seahorse assay plates. Compounds were loaded into a hydrated cartridge in the following order: Port A 1 µM Oligomycin, Port B 2µM FCCP, Port C 1 µM Rotenone/ Antimycin A, and Port D 20 µM Monensin. Measurements were determined by WAVE software. (**A-B**) Time-resolved plots showing (**A**) mitochondrial respiration (oxidative phosphorylation, OXPHOS) capacity as measured by oxygen consumption rate (OCR) and (**B**) glycolysis as measured by extracellular acidification rate (ECAR). (**C**) Stacked bar graph showing ATP production rates by glycolysis (J-ATP glyc) and mitochondrial respiration (J-ATP ox) of COMMA1D INSR cells under basal conditions. (**D**) Bar graph showing mitochondrial respiration ATP production rates (J-ATP ox) of COMMA1D INSR cells under maximal OXPHOS conditions (following FCCP injection). (**E**) Bar graph showing glycolytic ATP production rates (J-ATP glyc) of COMMA1D INSR cells under maximal glycolytic conditions (following Monensin injection). (**F**) Immunoblot of INSR and INSR KD reexpression in COMMA1D INSR (-/-) cells. ERK2 immunoblot is a loading control. (**G**) Bioenergetic plot representing the average glycolytic ATP production rate (J ATP-glyc; x-axis) plotted against the average mitochondrial respiratory ATP production rate (J ATP-ox; y-axis) of COMMA1D INSR (+/+), (-/-), and INSR (-/-) cells expressing the indicated constructs under basal, maximal OXPHOS (FCCP), and maximal glycolytic (Monensin) conditions. Time-resolved plots, bar graphs, and stacked bar graphs show mean with error bars representing SEM. * p < 0.05, **** p < 0.0001, n.s. = non-significant; Student’s t-test.

**Supplementary Table 1.**
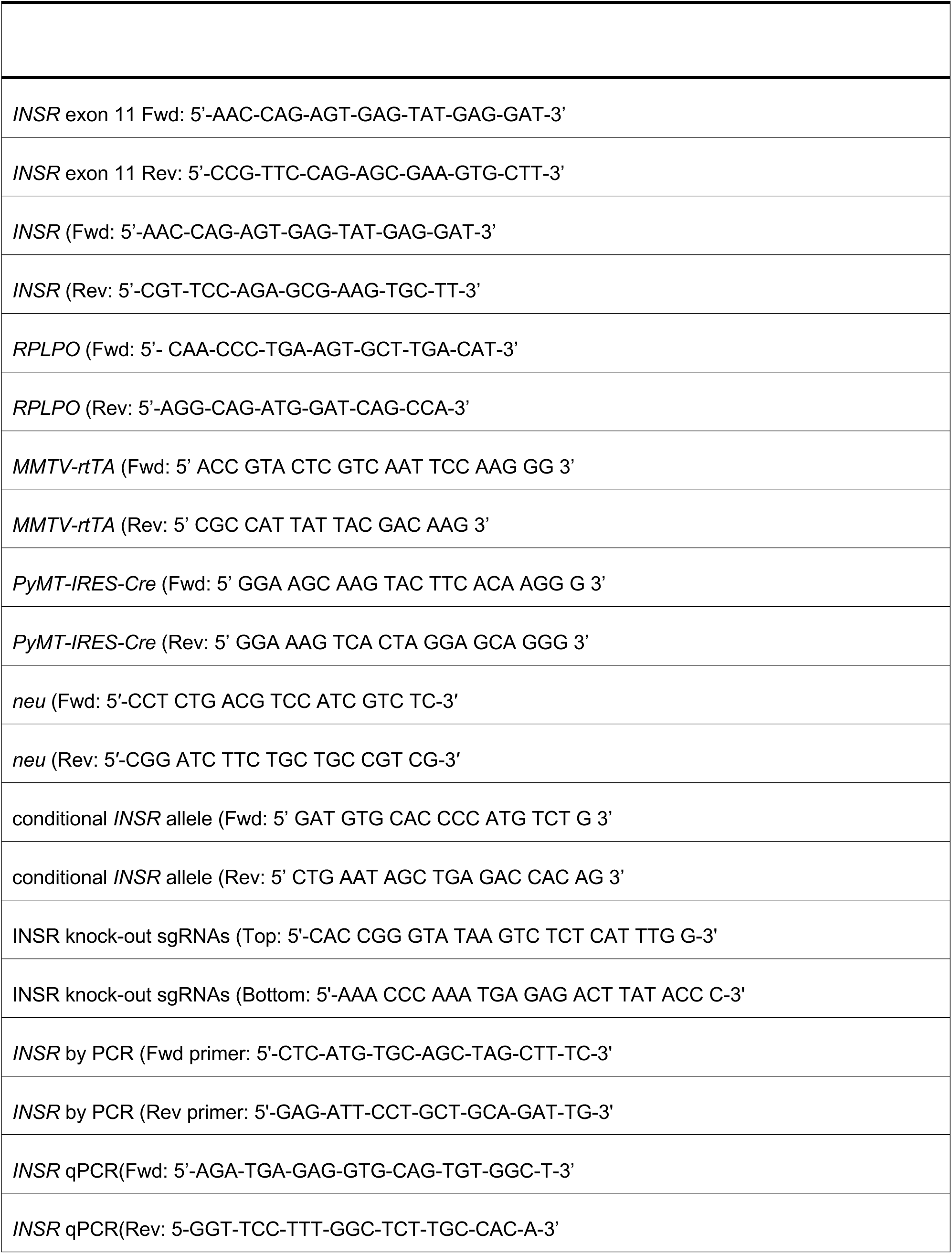
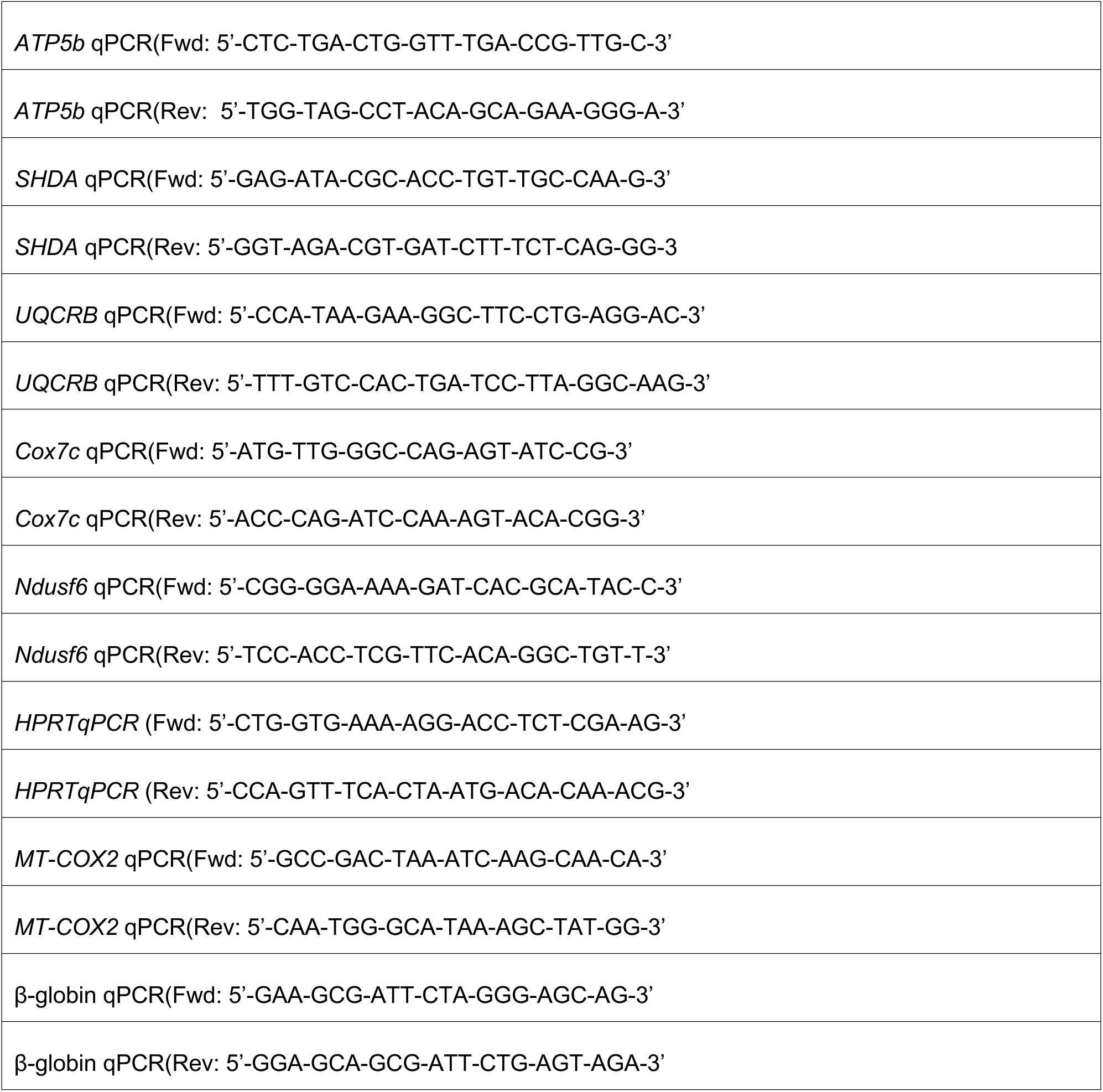
Oligonucleotides.

